# An immunogenic model of KRAS-mutant lung cancer for study of targeted therapy and immunotherapy combinations

**DOI:** 10.1101/2020.12.22.423126

**Authors:** Jesse Boumelha, Sophie de Carné Trécesson, Emily K. Law, Pablo Romero-Clavijo, Matthew A. Coelho, Kevin Ng, Edurne Mugarza, Christopher Moore, Sareena Rana, Deborah R. Caswell, Miguel Murillo, David C. Hancock, Prokopios P. Argyris, William L. Brown, Cameron Durfee, Lindsay K. Larson, Rachel I. Vogel, Alejandro Suárez-Bonnet, Simon L. Priestnall, Philip East, Sarah J. Ross, George Kassiotis, Miriam Molina-Arcas, Charles Swanton, Reuben Harris, Julian Downward

## Abstract

Mutations in oncogenes such as KRAS and EGFR cause a high proportion of lung cancers. Drugs targeting these proteins cause tumour regression but ultimately fail to cure these cancers, leading to intense interest in how best to combine them with other treatments, such as immunotherapies. However, preclinical systems for studying the interaction of lung tumours with the host immune system are inadequate, in part due to the low tumour mutational burden in genetically engineered mouse models. Here we set out to develop mouse models of mutant KRAS-driven lung cancer with an elevated tumour mutational burden by expressing the human DNA cytosine deaminase, APOBEC3B, to mimic the mutational signature seen in human lung cancer. This failed to substantially increase clonal tumour mutational burden and autochthonous tumours remained refractory to immunotherapy. However, by establishing clonal cell lines from these tumours we generated an immunogenic syngeneic transplantation model of KRAS mutant lung adenocarcinoma that was sensitive to immunotherapy. Unexpectedly, we found that anti-tumour immune responses were not directed against neoantigens but instead targeted derepressed endogenous retroviral antigens. The ability of KRAS^G12C^ inhibitors to cause regression of KRAS^G12C^-expressing versions of these tumours was markedly potentiated by the adaptive immune system, providing a unique opportunity for the study of combinations of targeted and immunotherapies in immune-hot lung cancer.

## INTRODUCTION

Non-small cell lung cancer (NSCLC) is the leading cause of cancer-related deaths worldwide^1^. With less than 20% of NSCLC patients surviving more than 5 years^2^, there is a pressing need for novel therapeutic strategies. Oncogenic mutations in KRAS, a member of the RAS family of small GTPases, occur in 20-30% of patients with NSCLC^3^ and drive multiple processes that promote tumour development. Despite much effort, targeted therapies that aim to directly inhibit signalling pathways downstream of KRAS have shown limited success in the clinic for NSCLC patients^4^. However, the recent emergence of immune checkpoint blockade (primarily anti-PD(L)-1 agents), which can reverse tumour-driven immune suppression and unleash powerful anti-tumour immune responses, has transformed the treatment of NSCLC, achieving durable responses in some patients^5^. Unfortunately, as seen in other tumour types, only a subset of patients respond to immune checkpoint blockade (ICB).

It has therefore become critical to further elucidate the molecular determinants that underpin the interaction between the tumour and the immune system. Increasing evidence suggests that tumour-cell-intrinsic oncogenic signalling, including KRAS signalling^6^, can hamper anti-tumour immune responses and there is considerable interest in using targeted therapies to broaden the clinical response to ICB. The recent development of KRAS^G12C^ inhibitors, which specifically target the most common mutant form of the protein in lung cancer, have shown that inhibiting KRAS-signalling in tumour cells promotes anti-tumour immune responses and synergises with anti-PD-1 therapy in an immune-competent model of colorectal cancer^7^.

Identifying rational therapeutic approaches to extend the clinical benefits of current ICBs in NSCLC requires the use of preclinical models that recapitulate the interactions between tumour cells and the immune system, which is not possible in conventional xenograft models lacking a functional immune system. Genetically engineered mouse models (GEMMs) have been extensively used to gain mechanistic insights into the biology of KRAS-mutant lung cancer and to assess the efficacy of novel therapeutics. Such models recapitulate key aspects of the human disease in an immune-competent setting, however, they fail to elicit strong anti-tumour immune responses^8,9^ and therefore have limited use for studying tumour-immune interactions. Genetically engineered mouse cancer models usually feature a small number of introduced strong driver mutations, sufficient for tumourigenesis, and acquire few additional mutations. Tumours arising from these models therefore have a low tumour mutational burden (TMB) compared to their human counterparts^10^, limiting the presentation of neoantigens to the adaptive immune system. This problem has been overcome by the forced expression of highly immunogenic antigens, such as ovalbumin^9,11^, but it is unclear whether the strong anti-tumour immune responses elicited by such foreign antigens reflect those in human cancers which occur towards less potent neoantigens and tumour-associated antigens.

To address this issue, we set out to generate a novel mouse model of KRAS driven lung adenocarcinoma with increased tumour mutation burden and potentially increased immunogenicity. Approaches used included the use of carcinogens and also the over-expression of a member of the APOBEC family of single-stranded DNA deaminases, which are responsible for inducing mutations in a range of cancers^12^. We were ultimately successful in generating a transplantable KRAS mutant lung cancer model that is partially sensitive to immunotherapy and shows a response to KRAS targeted agents that is clearly boosted by the adaptive immune system. This transplantable lung cancer model will be a valuable tool for studying strategies for combining targeted agents against the RAS pathway with immunotherapies.

## RESULTS

### Autochthonous KP lung tumours do not engage with the adaptive immune system

The introduction of adenovirus expressing Cre recombinase (AdCre) into the lungs of Kras^LSL-G12D/+^;p53^fl/fl^ (KP) mice leads to expression of oncogenic Kras^G12D^ and deletion of p53 in lung epithelial cells, resulting in the induction of lung adenocarcinoma^13^. This system represents one of the most widely used mouse models of lung cancer. To assess the immunogenicity of lung tumours arising in KP mice, we crossed them onto a *Rag2*^-/-^ background, which lacks mature T and B cells, and monitored tumour growth by micro-computed tomography (micro-CT) imaging. Adaptive immunity was unable to constrain the growth of KP tumours as they grew at similar rates in immune-competent (*Rag2*^+/−^) and immune-deficient (*Rag2*^-/-^) mice (Fig. 1A). Furthermore, immunohistochemistry staining revealed that tumours arising in immune-competent hosts lacked T cell infiltration (Fig. 1B). To assess whether an adaptive immune response could be generated against KP tumours, we treated tumour-bearing mice with a combination of anti-PD-L1 and anti-CTLA-4 (Fig. 1C). This combination therapy failed to delay tumour growth (Fig. 1D, E) and did not lead to an increase in the survival of tumour-bearing mice (Fig. 1F). It has previously been shown that MEK inhibition enhances anti-tumour immunity and synergises with anti-PD-L1 in KRAS-mutant CT26 colorectal tumours^14^. However, we found that the combination of anti-PD-L1 and trametinib failed to control KP tumour growth compared to trametinib alone (Supplementary Fig. 1A, B). These data suggest that the adaptive immune system is unable to recognise autochthonous KP tumours.

**Figure 1.**
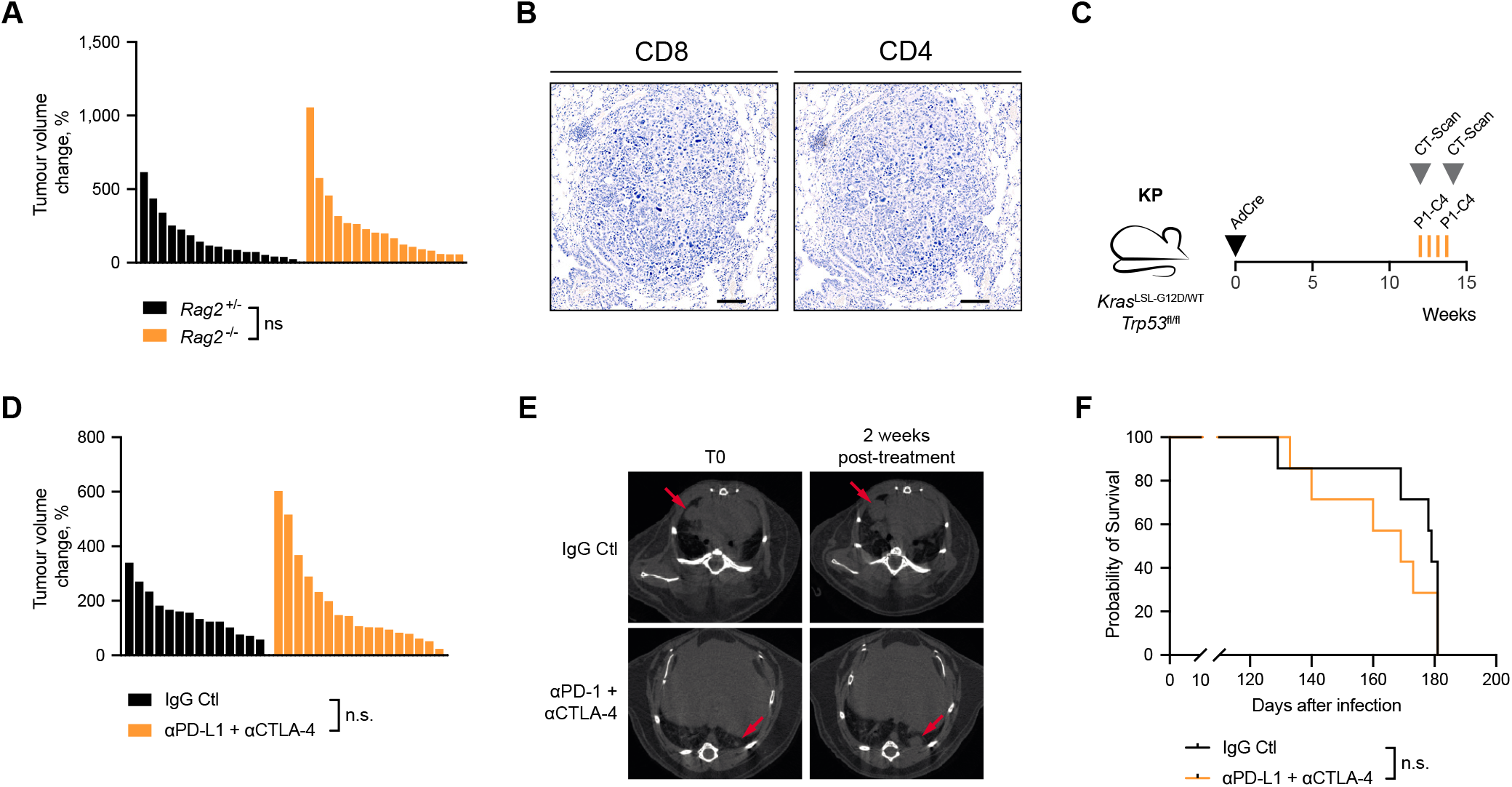
The KP mouse model of lung adenocarcinoma is not immunogenic. (A) Waterfall plot of tumour volume change over two weeks in KP;*Rag2*^+/−^(n=5) and KP;*Rag2*^-/-^ mice (n=4). Each bar represents volume change in a single tumour. One-way ANOVA, FDR 0.05; n.s. P>0.05. (B) Representative immunohistochemistry staining for CD4 and CD8 in KP tumours. Scale bar represents 100 μm. (C) Schematic of KP tumour induction and treatment schedule. Cre-expressing adenovirus (AdCre, 1×10^6^ pfu) was delivered intratracheally and mice were regularly scanned by micro-CT. 12 weeks after tumour initiation, tumour-bearing mice were treated three times (d0, d4 and d8) intraperitoneally with 10 mg/kg anti-PD-L1 and 5 mg/kg anti-CTLA-4 or corresponding isotype control (IgG Ctl). Tumour growth and survival were monitored until the experimental endpoints. (D) Waterfall plot of tumour volume change in KP-tumour-bearing mice treated as in (C) Mice were scanned 2 weeks after the pre-treatment scan, IgG Ctl (n=7) and anti-PD-L1 + anti-CTLA-4 (n=7). Each bar represents volume change in a single tumour. One-way ANOVA, FDR 0.05; n.s. P>0.05. (E) Representative micro-CT scans of mice treated as in (C). Red arrows indicate tumours. (F) Kaplan-Meier survival analysis of KP-tumour-bearing mouse survival treated as in (C), IgG Ctl (n=7) and anti-PD-L1 + anti-CT-LA-4 (n=7). Log-rank (Mantel-Cox) test; n.s. P>0.05

### Human APOBEC3B does not induce immunogenicity in the KP model

In comparison with human lung cancers seen in the clinic, KP mouse lung tumours exhibit very few mutations, which are necessary to generate neoantigens that can make tumour cells visible to the immune system^10^. APOBEC3B is a single-stranded DNA cytosine deaminase that induces C>T/G substitutions in several solid cancers^12,15,16^ and has been associated with intra-tumoural heterogeneity in lung adenocarcinoma^17,18^. Furthermore, analysis of lung adenocarcinoma (LUAD) samples from The Cancer Genome Atlas (TCGA) revealed that the mutational rate of non-synonymous APOBEC mutations was lower in comparison with other types of mutation, suggesting that non-synonymous mutations generated by APOBEC could be immunogenic and preferentially eliminated by the immune system (Fig. 2A).

**Figure 2.**
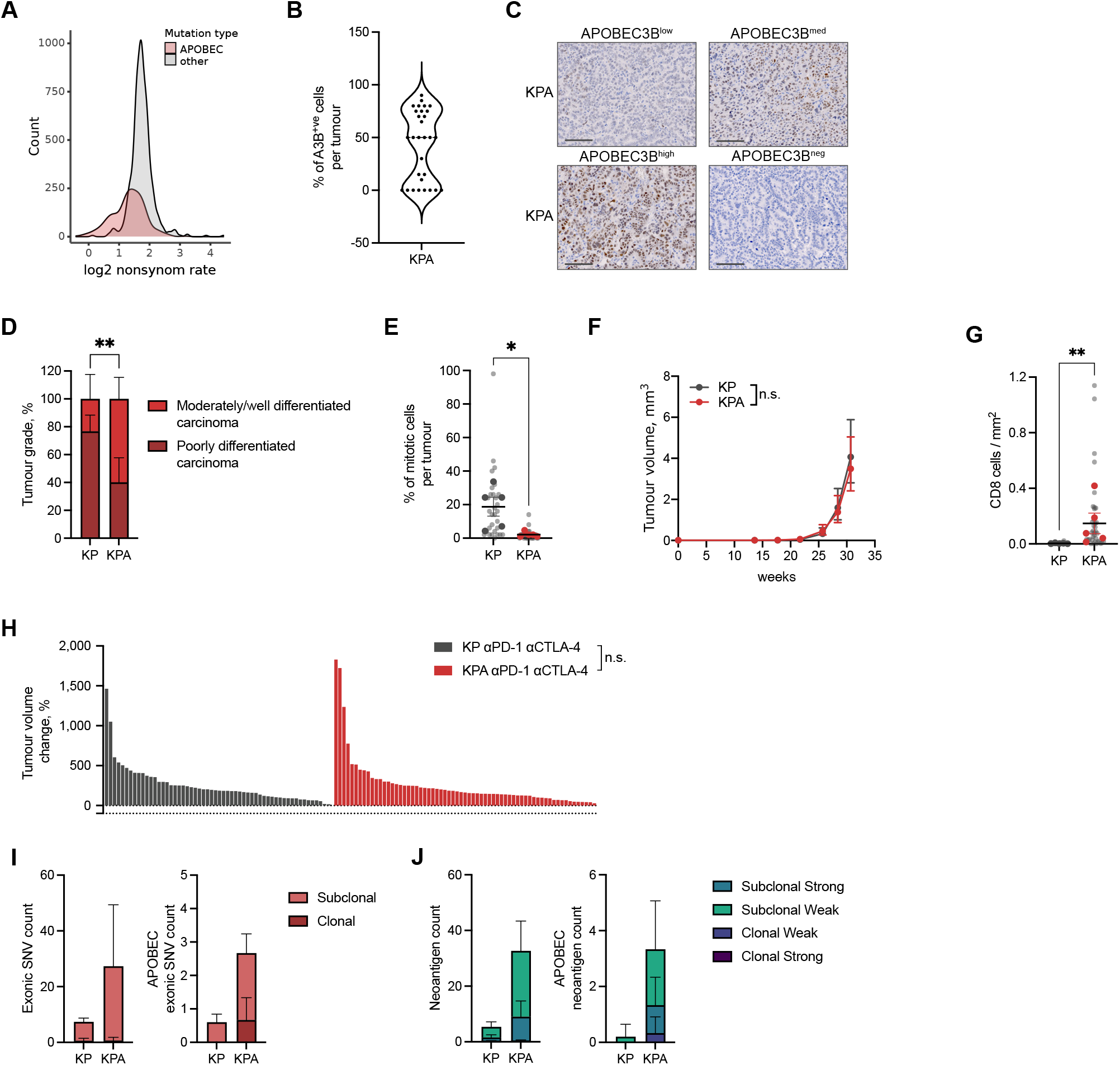
APOBEC3B expression in KP tumours is heterogeneous and deleterious for tumour progression. (A) Distribution of log2 non-synonymous/synonymous mutation ratio of APOBEC mutations or other types of mutation in LUAD (TCGA). (B) Distribution of A3Bi positive cells per lung tumour in the KPA model estimated by immunohistochemistry. (C) Immunohistochemistry of APOBEC3B staining in four KPA tumours showing different levels of APOBEC3B expression. Scale bar represents 100 μm. (D) Tumour grade proportion in the KP and KPA models. Percentage per model, upper and lower limit, Chi-square test p-value, n=5 mice per group; ** P≤0.01. (E) Percentage of mitotic cells in KP tumour and tumours expressing A3Bi in the KPA model estimated by histopathology. Light grey dots represent individual tumours. Mean per animal are represented in dark grey (KP) and red (KPA) dots. Mean per group, ±SEM and individual p-value, n=5 mice per group. One-way ANOVA of mean per group, FDR 0.05; * P≤0.05. (F) Tumour volume progression in KP (n=4 mice) and KPA (n=3 mice) models estimated by micro-CT scans. Geometric mean ± 95% CI. (G) Quantification of immunohistochemistry staining for CD8 in KP and KPA tumours. Light grey dots represent individual tumours. Mean per animal are represented in dark grey (KP) and red (KPA) dots. Mean per group, ±SEM and individual p-value, n=5 mice per group. One-way ANOVA of mean per group, FDR 0.05; ** P≤0.01. (H) Waterfall plot of tumour volume change in KP- and KPA-tumour-bearing mice treated with 200 μg of anti-PD-1 and 200 μg of anti-CTLA-4. Mice were scanned 2 weeks after the pre-treatment scan, KP (n=4) and KPA (n=7). Each bar represents volume change in a single tumour. One-way ANOVA, FDR 0.05; n.s. P>0.05. (I-J) Mean exonic SNV count ±SD (I) and mean neoantigen count ±SD (J) in KP (n=5 tumours) and KPA (n=3 tumours) broken down into clonal and subclonal. All neoantigens (left panels) and APOBEC-specific (T(C>T/G)) neoantigens (right panels).

We therefore decided to express human APOBEC3B in the KP model to increase the frequency of mutations in these tumours to promote the generation of neoantigens that could stimulate adaptive anti-tumour immune responses. We inserted a human *APOBEC3B* minigene (*A3Bi*) in the *Rosa26* locus under the control of a lox-STOP-lox cassette so that its expression is inducible upon exposure of cells to Cre recombinase (Supplementary Fig. 2A-J). A3Bi expression alone did not induce tumours and did not decrease the lifespan of the mice (Supplementary Fig. 2K-M). We crossed the mice with KP mice to generate *Kras*^G12D/+^;*Trp53*^fl/fl^;*Rosa26*^*A3Bi*^ (KPA) mice (Supplementary Fig. 3A). Following intratracheal AdCre delivery, A3Bi is initially expressed in the same cells as those that undergo *Kras*^G12D^ expression and *Trp53* deletion. As expected, we observed A3Bi protein expression in the nucleus of the tumour cells, however tumours contained different percentages of A3Bi-expressing cells (Fig. 2B and Supplementary Fig. 3B) as well as heterogenous levels of expression (Fig. 2C), suggesting a selection pressure against A3Bi expression during tumour growth. Consistent with this, KPA tumours were found to be of lower grade compared to KP tumours (Fig. 2D) and exhibited reduced tumour cell proliferation (Fig. 2E), as assessed by histopathology. However, KPA tumours grew at similar rates to KP tumours (Fig 2F). Interestingly, immunohistochemical analysis revealed that A3Bi expression was associated with moderate CD8^+^ T cell infiltration (Fig. 2G and Supplementary Fig 3C) which was confined to the periphery (Supplementary Fig 3D). In contrast, CD8^+^ T cells were entirely absent from KP tumours. To assess whether this increased CD8^+^T cell recruitment promoted immune-control of KPA tumours we crossed KPA mice onto a *Rag1*^-/-^ background to yield KPAR mice and evaluated tumour growth by micro-CT imaging in immune-competent KPA and immune-deficient KPAR animals. We observed no differences in tumour number or tumour growth in KPAR mice compared with KPA mice (Supplementary Fig. 3E, F). We also treated KP and KPA tumour-bearing mice with anti-PD-1 and anti-CTLA-4 and observed no differences between the two groups (Fig. 2H). We then performed whole-exome sequencing (WES) of KP and KPA tumours to assess whether A3Bi expression resulted in increased tumour mutational burden. We found that the total number of subclonal exonic SNVs was moderately increased in A3Bi-expressing tumours, however the majority of these were not typical A3Bi T(C>T/G) mutations (Fig. 2I). Consistent with this finding, KPA tumours possessed more predicted subclonal neoantigens, however few of these were likely to be directly due to APOBEC3B activity (Fig. 2J), perhaps indicating the possibility of indirect mechanisms linking APOBEC3B with the formation of new mutations.

Altogether, these findings suggest that A3Bi expression in the KP model of lung adenocarcinoma did not produce sufficient immunogenic mutations to elicit an adaptive immune response. This may be in part because of the heterogeneity of A3Bi expression in the tumours, the subclonal nature of any potential neoantigens, or alternatively due to insufficient numbers of mutations induced in this system.

### APOBEC3B increases subclonal mutations in urethane-induced tumours but does not promote immunogenicity

Murine KP tumours develop extremely rapidly, inducing a life-threatening tumour burden in about 14 to 18 weeks. We reasoned that the aggressive nature of the KP model did not allow sufficient time for APOBEC3B to induce mutations during tumour development, leading to only a few detectable SNVs with low allelic frequency. Carcinogen-induced tumours tend to be less aggressive than GEMMs and develop more slowly. To extend the length of tumour development, we exposed the mice to urethane before A3Bi expression. Urethane is a carcinogen which induces A>T/G substitutions and initiates lung tumours by inducing an activating mutation at codon Q61 in *Kras*^19,20^. To model A3Bi expression in carcinogen-induced tumours we initiated tumours with urethane in *Rosa26*^*A3Bi/CreER(t2)*^ mice (UrA3Bi) which when treated with tamoxifen express A3Bi in all tissues. Since APOBEC mutations are often late events in tumour evolution^17^, we delayed the induction of A3B by three weeks after the first injection of urethane (Supplementary Fig. 4A).

We confirmed the expression of A3Bi in the lungs after delivery of tamoxifen in UrA3Bi mice (Fig. 3A). A3Bi expression was downregulated in tumours compared with adjacent lung, consistent with what we observed in KPA tumours. Despite this, UrA3Bi tumours had more advanced histological grades compared with tumours induced by urethane alone (Fig. 3B). However, tumour growth and the number of tumours per animal were similar between urethane and UrA3Bi tumours (Fig. 3C and Supplementary Fig. 4B). To evaluate whether A3Bi expression promoted immunogenicity in urethane-induced tumours we treated UrA3Bi tumours with anti-PD-1 and anti-CTLA-4, however this combination failed to control tumour growth (Fig. 3D). We performed WES to assess whether A3Bi generated mutations in these tumours. As expected, urethane exposure generated a substantial number of clonal exonic SNVs (Fig. 3E). Similar to the KPA model, UrA3Bi tumours contained higher numbers of subclonal exonic SNVs (Fig. 3E) and predicted neoantigens (Fig. 3F) compared to urethane-induced tumours, however A3Bi expression failed to induce typical APOBEC T(C>T/G) mutations.

**Figure 3.**
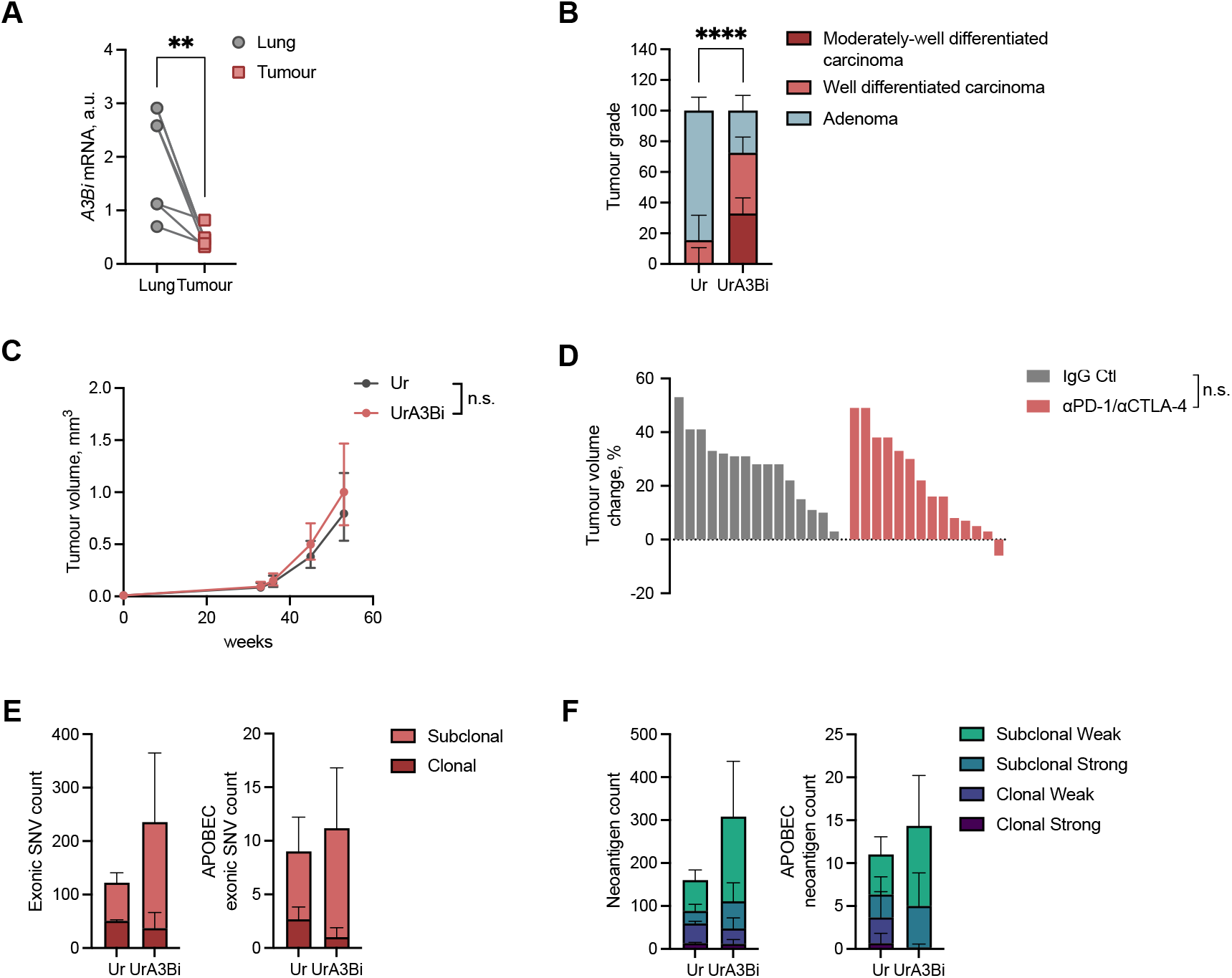
APOBEC3B expression increases mutations and tumour grade in p53-proficient autochthonous lung tumours. (A) Expression of A3Bi by qPCR in paired normal-adjacent tissue and tumour of UrA3Bi (n=7 tumours from 4 mice, squares), each symbol represents one tumour or adjacent tissue. Relative expression is normalised on the mean expression of *Sdha*, *Tbp* and *Actb*. Two-tailed paired t-test; ** P≤0.01 (B) Proportion of tumour grades evaluated from H&E staining of tumour-bearing lungs in Ur (n=8 mice) and UrA3Bi (n=8 mice) models. Chi-square test; **** P≤0.0001. (C) Tumour volume progression in Ur (n=9 mice) and UrA3Bi (n=10 mice) models estimated by micro-CT scans. Mean per model ±SEM two-way ANOVA =, FDR 0,05; n.s. P>0.05 (D) Waterfall plot of tumour volume change in UrA3Bi-CreER mice. Tumour-bearing mice were scanned and treated four times (d0, d3, d7 and d10) with 200 μg of anti-PD-1 and 200 μg of anti-CTLA-4 (n=2 mice) or corresponding isotype control (n=2 mice). mice were scanned 2 weeks after the pre-treatment scan. Each bar represents volume change in a single tumour. Two-way ANOVA; n.s.= P>0.05 (E) Mean exonic SNVs count ±SD in Ur (n=3 tumours) and UrA3Bi (n=6 tumours) broken down into clonal and subclonal. All exonic SNVs (left panel) and APOBEC-specific (T(C>T/G)) (right panel). (F) Mean neoantigen count ±SD in Ur (n=3 tumours) and UrA3Bi (n=6 tumours) broken down into weak and strong binders from clonal and subclonal SNVs. All neoantigens (left panel) and APOBEC-specific (T(C>T/G)) neoantigens (right panel).

To summarise, as with the KP model, APOBEC3B expression failed to induce immunogenicity in carcinogen-induced models of lung cancer.

### Establishment of immunogenic clonal cell lines from KPAR tumours

We hypothesised that the lack of immunogenicity in APOBEC3B-expressing autochthonous tumours might be due to the subclonality of mutations, which have been shown to be less effective at generating effective adaptive immune responses^21,22^. We therefore established cell lines from these models which were subsequently single-cell cloned to increase the frequency of clonal neoantigens. Cell lines established from urethane-induced tumours grew poorly *in vitro* and failed to grow when transplanted into mice (data not shown), probably as urethane-induced tumours are typically very low grade, often only possess a mutation in Kras (Q61R) and lack any further oncogenic alterations. However, a number of cell lines were readily established from KPA and KPAR autochthonous tumours and single-cell cloned. We were unable to detect APOBEC3B mRNA expression in any of the KPAR cell lines (Fig. 4A), suggesting that expression of the transgene was downregulated during tumour growth, consistent with what we observed by immunohistochemistry (Fig. 2B). In contrast, APOBEC3B mRNA expression was detected in the KPA cell lines and therefore we decided not to further characterise them as the expression of a human recombinant protein could affect the growth of transplanted tumours in immune-competent hosts.

**Figure 4.**
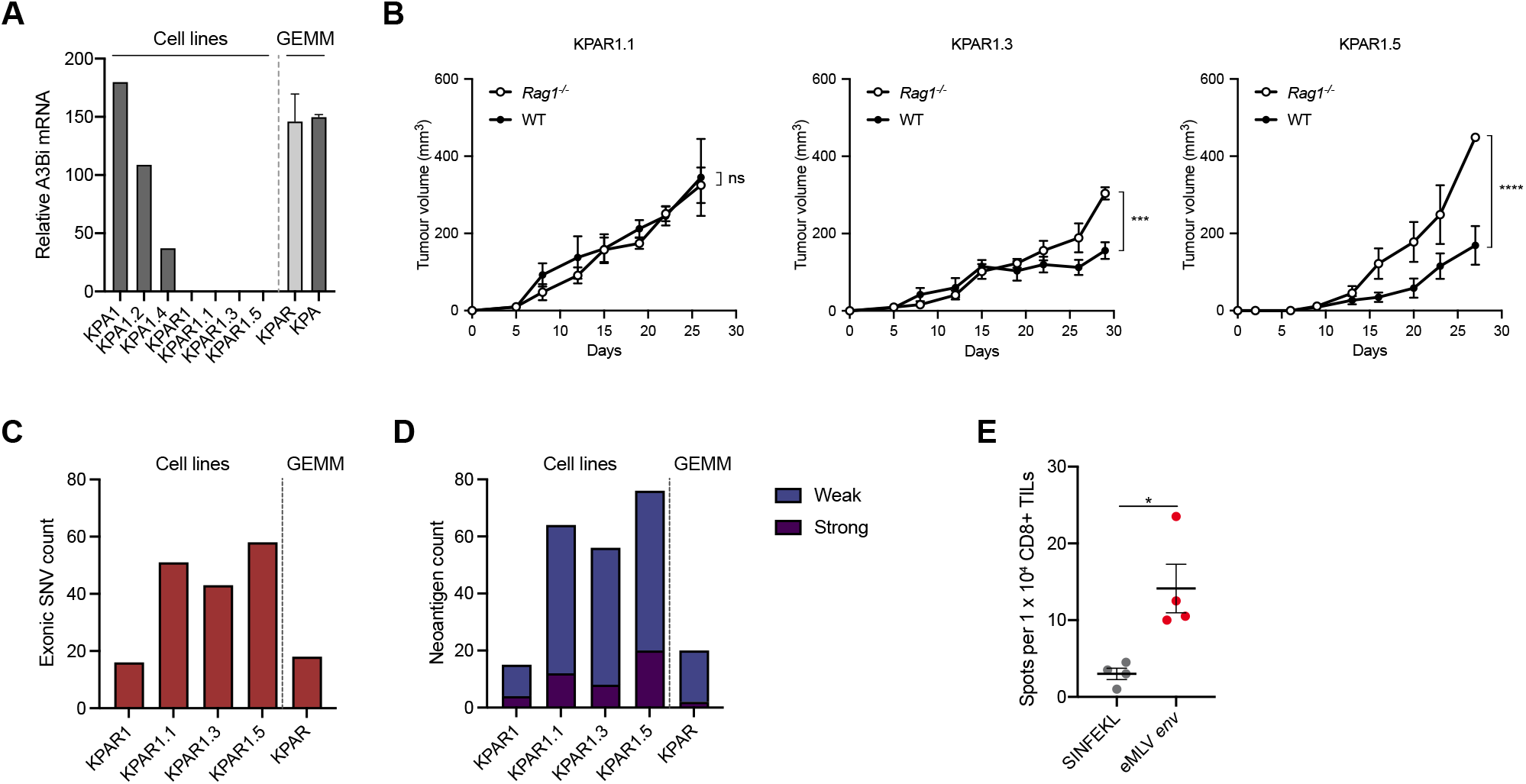
Generation of a novel immunogenic cell line KPAR1.3. (A) mRNA expression by qPCR of A3Bi in autochthonous KPA and KPAR tumours, KPA and KPAR parental cells and KPA and KPAR sub-clones. All values are normalised to the mean expression of *Sdha*, *Tbp* and *Hsp90ab1*. (B) Growth of KPAR cells transplanted subcutaneously into syngeneic immune-competent and *Rag1*^-/-^ mice. Data are mean tumour volumes ± SEM, KPAR1.3 and KPAR1.5 n=5 mice per group and KPAR1.1 n=4 mice per group. Two-way ANOVA, *** P≤0.001, **** P≤0.0001, n.s. = P>0.05 (C) Frequency of exonic mutations in an autochthonous KPAR tumour, the KPAR parental cell lines and the KPAR1.1, KPAR1.3 and KPAR1.5 single-cell clones, estimated par whole-exome sequencing. (D) Frequency of predicted neoantigens identified using NetMHC4.0. Peptides with a rank threshold of <2 or <0.5 were designated as weak or strong MHC-I binders, respectively. (E) IFNγ ELISPOT analysis of CD8^+^ TILs isolated from KPAR1.3 subcutaneous tumours and pulsed with indicated peptides. SINFE-KL was used as a negative control. Data are mean ± SEM, n=4 mice per group. Unpaired, two-tailed Student’s t-test; * P≤0.05,

The immunogenicity of different single-cell KPAR clones was assessed by comparing the growth of cells subcutaneously transplanted into syngeneic immune-competent and immune-deficient (*Rag1*^-/-^) mice. KPAR1.1 cells grew similarly when injected into immune-competent and *Rag1*^-/-^ mice whilst two other clones, KPAR1.3 and KPAR1.5, grew more slowly in immune-competent mice compared to *Rag1*^-/-^ mice (Fig. 4B).

We carried out WES to assess the mutational burden of the KPAR clonal cell lines, the parental polyclonal cell line (KPAR1) and another autochthonous KPAR tumour taken from the same mouse. Single-cell cloning moderately increased the frequency of detectable mutations (Fig. 4C). All single-cell clones contained more mutations compared with the parental cell line or KP tumours (Fig. 2I). However, the number of mutations in all clonal cell lines was still very low compared to other transplantable syngeneic cancer cell lines, and immunogenic KPAR1.3 cells did not possess more predicted neoantigens than non-immunogenic KPAR1.1 cells (Fig. 4D, Supplementary Table S1). Notably, very few mutations were typical APOBEC T(C>T/G) mutations (Supplementary Fig. 5A). Furthermore, we were unable to detect antigen-specific CD8^+^ T cells against any of the predicted neoantigens when pulsing tumour-infiltrating lymphocytes (TILs) isolated from KPAR1.3 tumours in an IFNγ enzyme-linked immune absorbent spot (ELISpot) assay (Supplementary Table S1).

Given that A3Bi failed to induce any immunogenic mutations in the KPAR1.3 cell line we asked whether the immunogenicity of this cell line may be due to another source of antigens. One major class of tumour-associated antigens consists of endogenous retroviral proteins that are often derepressed in established mouse cancer cell lines^23^ and have also been shown to drive anti-tumour immune responses in human cancer^24^. Interestingly, ELISPOT analysis revealed that the major MHC-I restricted epitope arising from the envelope glycoprotein (*env*) of *Emv2*, the endogenous ecotropic murine leukaemia retrovirus (eMLV) in C57BL/6J mice, induced IFNγ secretion from CD8^+^ KPAR1.3 TILs (Fig. 4E). Consistent with this, immunogenic KPAR1.3 and KPAR1.5 clones expressed significantly higher levels of endogenous MLV envelope proteins compared to the non-immunogenic KPAR1.1 cell line (Supplementary Fig. 5B).

Together these results indicate that the immunogenicity of the KPAR1.3 cell line was not due to tumour mutational burden, but elevated expression of endogenous retroviral antigens that stimulate endogenous CD8^+^ T cell responses.

### KPAR tumours generate an adaptive immune response

Although immunogenicity of the KPAR1.3 cell line was not due to neoantigens generated by non-synonymous mutations as we initially hypothesised, the novelty of an immunogenic transplantable murine lung cancer cell line warranted further characterisation. We therefore used flow cytometry to characterise the tumour microenvironment of orthotopic lung tumours established from immunogenic KPAR1.3 cells (from now on referred to as KPAR) and non-immunogenic KPB6 cells, derived from the original KP GEMM on a C57BL/6J background. Notably, KP tumours have very few predicted neoantigens (Fig. 2J) and KPB6 cells display significantly reduced surface expression of endogenous MLV envelope proteins compared to KPAR cells (Supplementary Fig. 5C). The immune compartment of both tumour models differed significantly compared to normal lung with a large increase in the proportion of myeloid cells, consisting primarily of interstitial macrophages, and exclusion of B cells and NK cells (Fig. 5A). KPB6 tumours contained significantly more myeloid cells than KPAR tumours, primarily due to an increased proportion of neutrophils (Supplementary Fig. 6A). Conversely, KPAR tumours showed significantly higher levels of T cell infiltration, which was a result of increased CD8^+^ and CD4^+^ T cells and Tregs, as well as increased NK cell infiltration (Fig. 5B). Immunohistochemistry staining confirmed that KPAR tumours were more infiltrated with CD8^+^ T cells (Supplementary Fig. 6B). T cells infiltrating KPAR tumours were also more activated, with a higher proportion of effector memory CD8^+^ and CD4^+^ T cells (Fig. 5C) and increased expression of the activation marker CD44 on both CD8^+^ and CD4^+^ T cells (Supplementary Fig. 6C, D) as well as the early activation marker CD69 on CD8^+^ T cells (Supplementary Fig. 6E). Both CD8^+^ and CD4^+^ T cells also showed increased expression of the immune checkpoint molecules PD-1, LAG-3 and TIM-3 in KPAR tumours (Fig. 5D). Furthermore, KPAR tumours also contained a significant proportion of PD-1/LAG-3 double-positive CD8^+^ T cells which were completely absent in KPB6 tumours (Fig. 5E). There was also an increased proportion of PD-L1^+^ myeloid cells in KPAR tumours, indicative of a T-cell inflamed tumour microenvironment (Fig. 5F).

**Figure 5.**
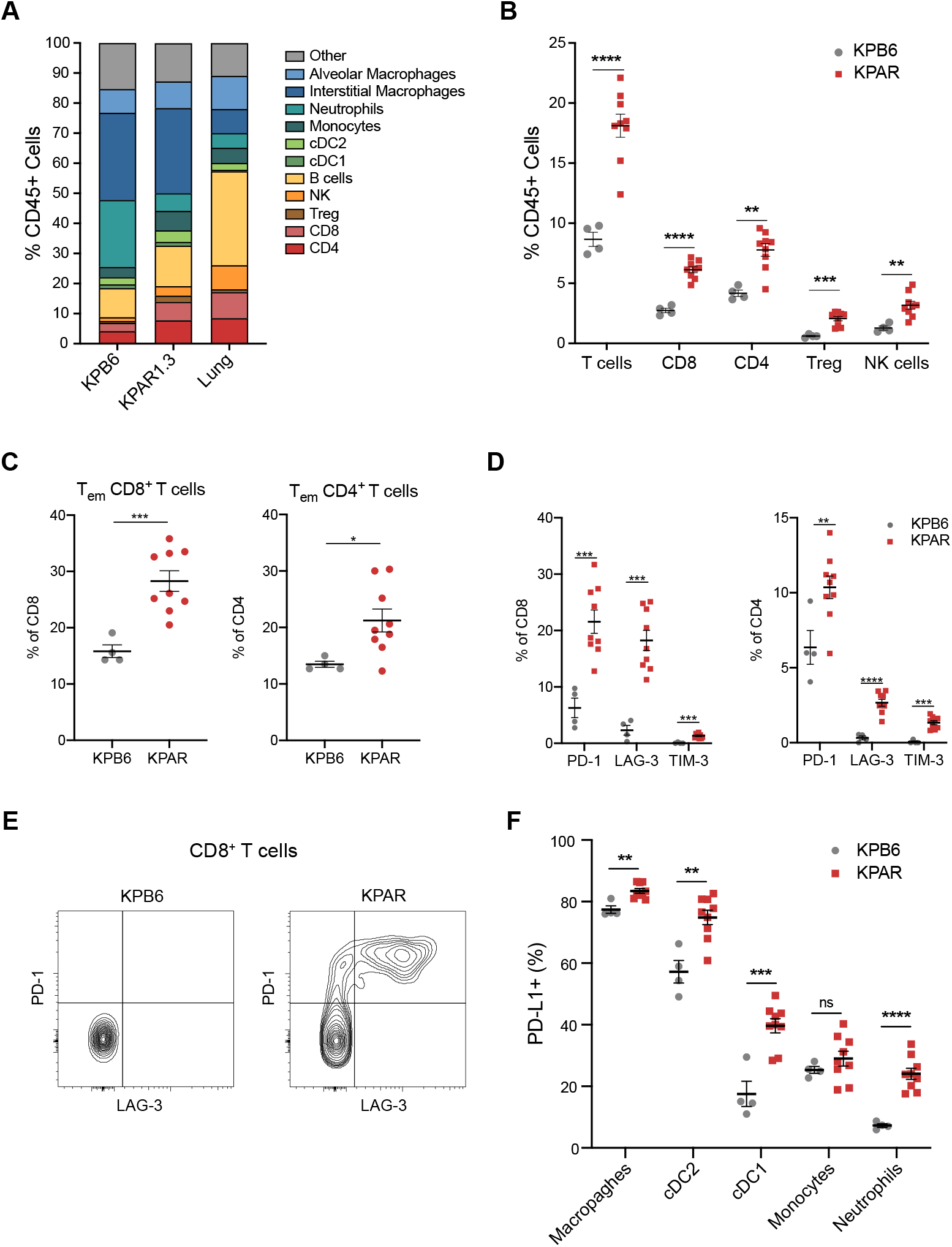
KPAR orthotopic tumours generate an adaptive immune response. (A) Immune profile of KPARand KPB6 orthotopic tumours compared to normal lung, assessed by flow cytometry. (B) Frequency of tumour-infiltrating T cell populations and NK cells (CD11b-CD49b^+^Nkp46^+^). (C) Percentage of effector memory (CD44^+^CD62L^−^) CD8^+^ (left) and CD4^+^ (right) T cells. (D) Quantification of PD-1, LAG-3 and TIM-3 expression on CD8^+^ (left) and CD4^+^ (right) T cells. (E) Representative plot of PD-1 and LAG-3 expression on CD8^+^ T cells. (F) Frequency of PDL1^+^ macrophages (CD11b^+^MHCII^+^CD64^+^), cDC2 (CD11b^+^MHCII^+^CD24^+^), cDC1 (CD11c^+^CD64-CD24^+^CD103^+^), monocytes (CD11b^+^Ly6C^+^) and neutrophils (CD11b^+^Ly6G^+^). In (A)-(D) and (F), data are mean ± SEM, n=4 mice (KPB6) or 9 mice (KPAR), symbols represent pooled tumours from individual mice. In (A)-(F) tumours were analysed 21 days after transplantation. Unpaired, two-tailed Student’s t-test; ns P>0.05, * P≤0.05, ** P≤0.01, *** P≤0.001, **** P≤0.0001.

Taken together, these data demonstrate that orthotopic KPAR tumours generated an adaptive anti-tumour immune response which was absent in orthotopic KPB6 tumours.

### KPAR tumours are responsive to ICB

Given that the growth of KPAR tumours were partially restrained by adaptive immunity and orthotopic tumours were highly infiltrated with activated immune cells, we next tested the sensitivity of the model to ICB. Mice bearing subcutaneous KPAR tumours were treated with anti-PD-1, anti-CTLA-4 or a combination of both. Anti-CTLA-4 alone, or in combination with anti-PD-1, led to tumour regression in all mice, whilst anti-PD-1 alone failed to affect tumour growth. (Fig. 6A, and Supplementary Fig. 7A). Furthermore, anti-CTLA-4 or the combination of anti-CTLA-4 and anti-PD-1 resulted in long-term durable regression for up to one year in 33% and 50% of mice, respectively (Fig. 6B). All treated mice that had rejected the primary tumour subsequently rejected a secondary tumour when re-challenged with KPAR cells on the opposite flank (Fig. 6B), demonstrating the establishment of immunological memory. Furthermore, we observed significantly more IFNγ spots by ELISpot analysis when CD8^+^ T cells isolated from subcutaneous KPAR tumours treated with anti-CTLA-4 were pulsed with the eMLV envelope peptide compared to isotype control, indicating that eMLV-specific T cells expand in response to immunotherapy (Supplementary Fig. 7B).

**Figure 6.**
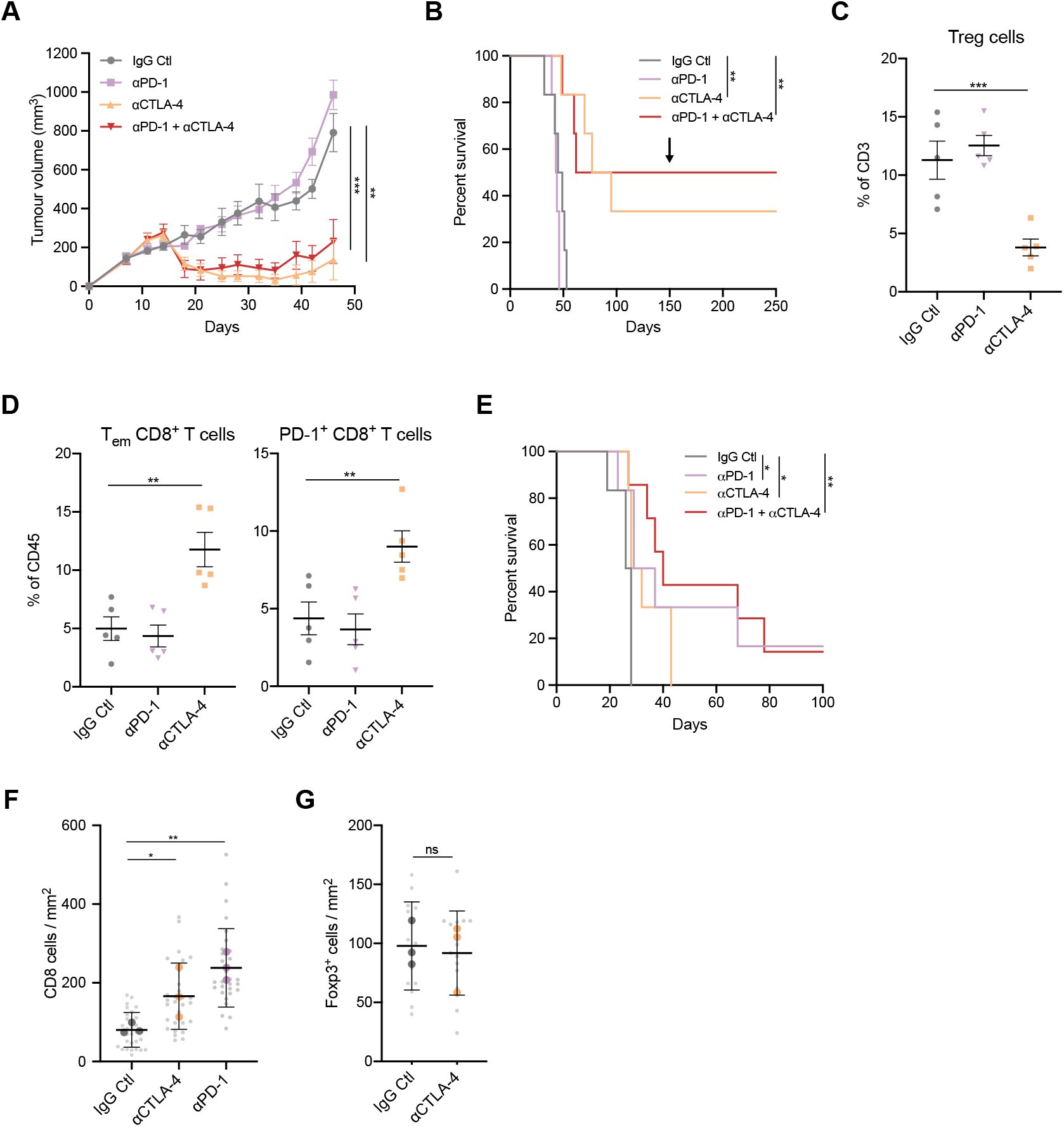
Subcutaneous and orthotopic KPAR tumours are responsive to ICB. (A) Growth of KPAR subcutaneous tumours from mice treated intraperitoneally with 200μg anti-PD-1 and/or 200μg anti-CTLA-4 or corresponding isotype control (IgG Ctl) on day 10, 14, 17 and 21. Data are mean tumour volumes ± SEM, n=6 mice per group. Two-way ANOVA; ** P≤0.01, *** P≤0.001. (B) Kaplan-Meier survival of mice from (A). The black arrow indicates the time at which mice that previously rejected the primary tumour were re-challenged on the opposite flank. Log-rank (Mantel-Cox) test; ** P≤0.01. (C and D) Flow cytometry analysis of the frequency of Foxp3^+^ Tregs (C), effector memory (CD44^+^CD62L^-^) CD8^+^ T cells (D, left) and PD-1^+^ CD8^+^ T cells (D, right) in subcutaneous tumours after treatment as in (A). Treatment was on day 10, 14 and 17 and mice were culled on day 18. Data are mean ± SEM, n=5 mice per group. One-way ANOVA, FDR 0.05; ** P≤0.01, *** P≤0.001. (E) Kaplan-Meier survival of mice treated intraperitoneally with 200μg anti-PD-1 and/or 200μg anti-CTLA-4 or corresponding isotype control (IgG Ctl) after orthotopic transplantation of KPAR cells. Treatment was initiated once tumours were detectable by micro-CT and were administered twice weekly for a maximum of 3 weeks. IgG Ctl (n=6), anti-PD1 (n=6), anti-CTLA4 (n=6) and anti-PD-1 + anti-CTLA-4 (n=7). Log-rank (Mantel-Cox) test; * P≤0.05, ** P≤0.01. (F and G) Quantification of immunohistochemistry staining for CD8 (F) and Foxp3 (G) in orthotopic KPAR lung tumours after treatment as in (E). Data are mean (large symbols) ± SEM, n=3 mice per group, small symbols represent individual tumours. One-way ANOVA, FDR 0.05; ns P>0.05, * P≤0.05, ** P≤0.01.

Flow cytometry analysis of subcutaneous tumours treated with anti-CTLA-4 or anti-PD-1 demonstrated that only anti-CTLA-4 treatment effectively depleted Foxp3^+^ regulatory T cells (Tregs) (Fig. 6C), resulting in an increase in the ratio of CD8^+^ and CD4^+^ effector T cells to Tregs (Supplementary Fig. 7C, D), as previously reported^25^. Anti-CTLA-4 treatment also led to an increase in the frequency of effector memory and PD-1^+^ CD8^+^ T cells in tumours (Fig. 6D). To assess whether the sensitivity to ICB was dependent on the anatomic site of tumour growth, as previously shown^26^, we also treated orthotopic KPAR lung tumours with anti-PD-1, anti-CTLA-4 or a combination of both. KPAR cells were injected intravenously into mice which were subsequently treated once lung tumours were detected by micro-CT. In contrast to subcutaneous tumours, orthotopic tumours responded to both anti-CTLA-4 and anti-PD-1 monotherapies, resulting in a significant increase in the survival of tumour-bearing mice (Fig. 6E). However, the response to anti-PD1 was substantially greater, resulting in long-term responses in a subset of mice as a monotherapy or in combination with anti-CTLA-4. Immunohistochemistry staining demonstrated that both anti-PD-1 and anti-CTLA-4 therapy increased the infiltration of CD8^+^ T cells into the tumour (Fig. 6F, and Supplementary Fig. 7E), however this increase was greater in anti-PD-1 treated mice. In contrast to subcutaneous tumours, anti-CTLA-4 treatment failed to deplete Foxp3^+^ Tregs in orthotopic tumours (Fig. 6G, and Supplementary Fig. 7F). Furthermore, the majority of CD4^+^ T cells in subcutaneous tumours were Tregs whilst in orthotopic tumours CD4^+^ effector T cells were more abundant (Supplementary Fig. 7G).

To summarise, KPAR1.3 tumours were sensitive to anti-PD-1 or anti-CTLA-4 immune checkpoint blockade therapy, the response to which was dependent on the site of tumour growth.

### Generation of KPAR^G12C^ cells to assess the immunomodulatory properties of KRAS^G12C^ inhibitors

The recently developed class of KRAS^G12C^ inhibitors has been shown to promote anti-tumour immune responses in the immunogenic CT26^G12C^ model of colorectal cancer^7^.

To test the effect of KRAS^G12C^ inhibitors in the KPAR lung cancer model we used prime-editing technology to generate the KPAR^G12C^ cell line. WES revealed that KPAR cells were homozygous for KRAS^G12D^ so we edited both alleles to KRAS^G12C^ (Supplementary Fig. 8A). Cell-viability assays demonstrated that KPAR^G12C^ cells showed impaired viability in response to treatment with AZ-8037, a recently described KRAS^G12C^ inhibitor ^27^ (Supplementary Fig. 8B). Furthermore, immunoblotting revealed that AZ-8037 inhibited pERK in KPAR^G12C^ cells (Supplementary Fig. 8C). To assess whether KRAS^G12C^ inhibition could stimulate anti-tumour immunity *in vivo* we tested the response of KPAR^G12C^ subcutaneous tumours to AZ-8037 in both immune-competent and immune-deficient (*Rag1*^-/-^) mice. Vehicle-treated KPAR^G12C^ tumours grew slower in immune-competent mice compared to *Rag1*^-/-^ mice, similarly to what we observed with the parental KPAR tumours (Fig. 7A). AZ-8037 treatment caused marked tumour regression in both immune-competent and *Rag1*^-/-^ mice, however the response was much more durable in immune-competent mice as all tumours remained responsive during the duration of treatment whilst tumours in *Rag1*^-/-^ mice began to grow back before termination of treatment (Fig. 7A). Furthermore, after the treatment was terminated one of the six treated mice showed a durable cure (Fig. 7B). We also used CRISPR technology to edit the KPB6 cell line, which harbours a wildtype KRAS and KRAS^G12D^ allele, to generate the KPB6^G12C^ cell line which lost the wildtype allele by indel generation and contained a KRAS^G12C^ allele (Supplementary Fig. 8D). Cell-viability assays and immunoblotting demonstrated that KPB6^G12C^ cells were sensitive to AZ-8037 (Supplementary Fig. 8E, F). In contrast to KPAR^G12C^ tumours, the response of KPB6^G12C^ tumours to AZ-8037 was comparable in immune-competent and *Rag1*^-/-^ mice (Fig. 7C), with tumours beginning to lose responsiveness before treatment ended, and then growing back rapidly after the cessation of treatment with no long-term responses achieved (Fig. 7D). Given that adaptive immunity contributes to the efficacy of KRAS^G12C^ inhibition in KPAR tumours, we next wanted to assess the effects of KRAS^G12C^ inhibition on the tumour microenvironment. qPCR analysis of orthotopic KPAR^G12C^ tumours revealed that KRAS^G12C^ inhibition induced a pro-inflammatory microenvironment with increased antigen presentation, cytokine production, interferon signalling, immune cell infiltration and T cell activation (Fig 7E).

**Figure 7.**
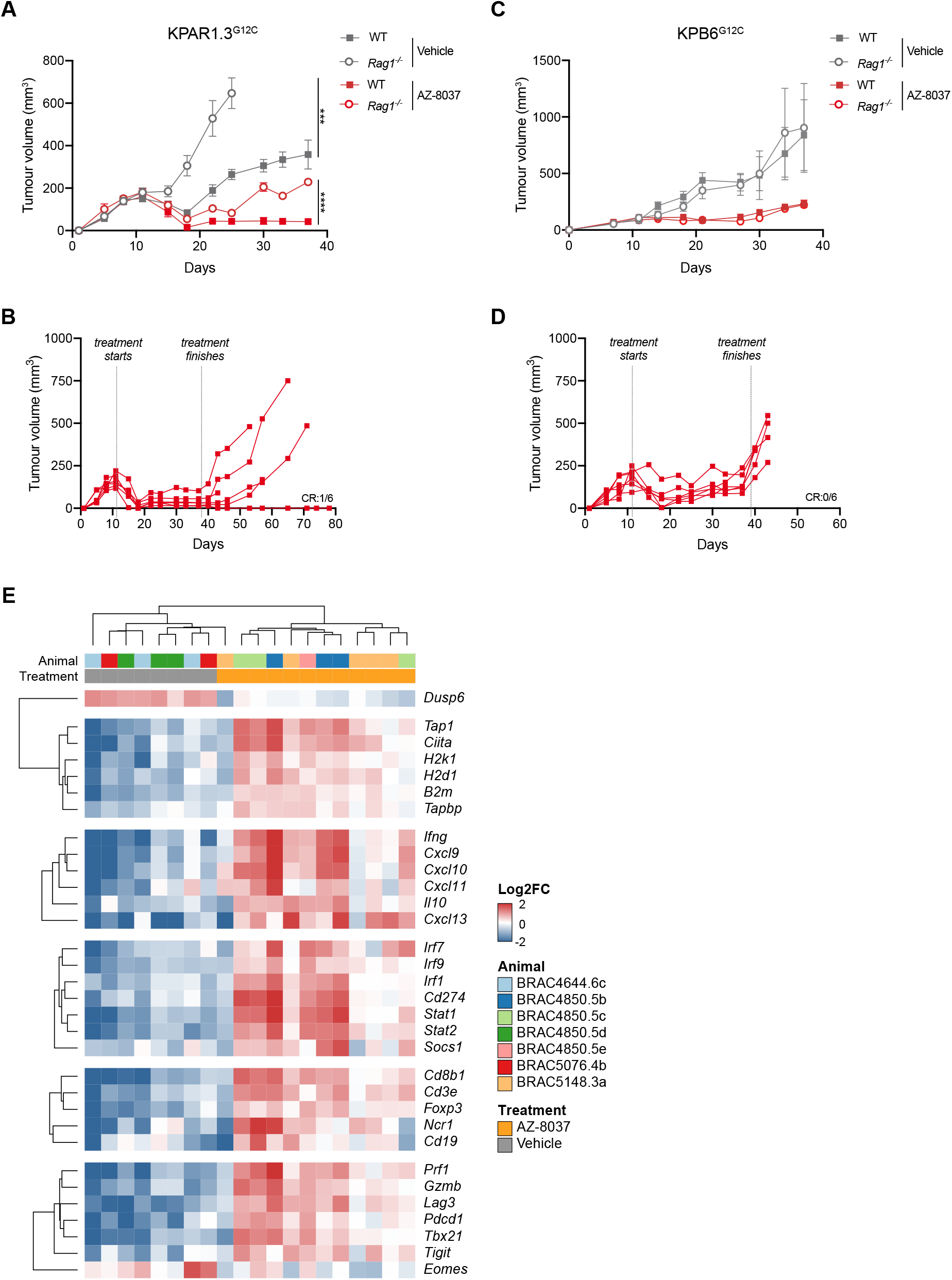
The efficacy of KRAS-G12C inhibition in vivo is greater in immune-competent mice. (A and B) Mean ± SEM (A) and individual (B) KPAR1.3G12C tumour volumes in immune-competent and *Rag1*^-/-^ mice treated with vehicle or AZ-8037 (100 mg/kg daily oral gavage). n=6 mice per group. Two-way ANOVA; *** P≤0.001, **** P≤0.0001. (C and D) Mean ± SEM (C) and individual (D) KPB6G12C tumour volumes in immune-competent and *Rag1*^-/-^ mice treated as in (C). n=6 mice per group. (E) Heatmap showing mRNA expression from qPCR of KPAR1.3G12C tumours treated for 7 days with 100 mg/kg AZ-8037. Gene expression is scaled across all tumours. Only genes with a significant mean difference between AZ-8037 and vehicle groups (one-way ANOVA, FDR<0.05) are shown.

These results suggest the efficacy of KRAS^G12C^ inhibition in the immunogenic KPAR model was partially due to the generation of an adaptive anti-tumour immune response which resulted in durable regressions in immune-competent hosts.

## DISCUSSION

There is a need for improved models of lung cancer that are immunogenic to enable us to better understand the interplay between the tumour and the immune system and assess the efficacy of novel therapeutic interventions. We and others have tried several approaches to make lung cancer GEMMs more immunogenic. We treated KP mice with carcinogens, expressed APOBEC3B in p53-deleted urethane-induced tumours, and – based on the assumption that chromosome rearrangement could lead to neoantigens – also expressed Mad2, a spindle checkpoint protein associated with aneuploidy, in KP tumours^28^. However, none of these strategies generated immunogenic tumours that grew differentially in immune-competent and immune-deficient backgrounds, induced T cell responses or responded to immune checkpoint blockade (data presented here and unpublished data). In this study, APOBEC3B expression only moderately increased the tumour mutational burden in KP and urethane-induced lung tumours and was not sufficient to make these tumours immunogenic. The lack of substantial numbers of APOBEC3B induced mutations in these models was potentially a consequence of a detrimental impact of APOBEC3B expression during early stages of tumour development, which is reflected by the downregulation of A3Bi expression in both KP and urethane-induced lung tumours. The development of a KP model with a temporal regulation of APOBEC3B could help addressing this limitation. Furthermore, APOBEC3B-induced mutations were mainly subclonal in the autochthonous tumours, as described in patients^18^. The subclonality of many mutations in these models and the heterogenous A3Bi expression we observed in tumours could explain the absence of effective anti-tumour immune responses, since sensitivity to ICB correlates with clonal neoantigen burden in NSCLC^29^. However, despite containing many more clonal exonic mutations than A3Bi-expressing KP tumours, urethane-induced lung tumours were also refractory to ICB. The long latency of urethane-induce lung tumours may provide ample time for the elimination of immunogenic clones. Alternatively, a higher number of somatic mutations than achieved here may be required. Indeed, peptide screens of human tumour samples have revealed that only a minority of mutations result in neoantigens that are recognised by TILs^30^. Together these results highlight the limitations of autochthonous models of lung cancer which fail to induce anti-tumour immune responses.

Given the limitations of autochthonous models, most preclinical studies of immunotherapy utilise transplantable syngeneic cell lines. The most commonly used mouse cancer cell lines used for syngeneic transplantation that are sensitive to immunotherapy are the colorectal carcinoma cell line CT26 and renal cancer cell line RENCA, with many other commonly used lines such as MC38 colorectal cancer, B16-F10 melanoma and 4T1 breast cancer being largely refractory to immune checkpoint blockade^31^. Promoting DNA damage pathways in cells such as CT26 by deleting MLH1 DNA mismatch repair gene further increases tumour mutational burden and response to immunotherapy^32^. The most commonly used murine lung cancer cell line for orthotopic preclinical studies is the 3LL cell line, also referred to as LL/2 or LLC1, and derivative variants, which originate from a spontaneous Lewis lung carcinoma tumour in a C57/BL6 mouse that has been serially passaged in immune-competent mice, leading to a highly immune evasive phenotype^31^. It has activating mutations in both KRAS and NRAS^33^, however these tumours are refractory to ICB^34^, largely due to their ability to generate a very immunosuppressive tumour microenvironment rather than a lack of tumour neoantigens, and therefore do not make a suitable model for studying the response to novel therapy combinations in an immune-hot tumour microenvironment context. Another transplantable mouse lung cancer cell line, CMT-167, has been characterised as KRAS^G12V^ mutant and found to be responsive to immunotherapy^26^.

In this study we established the KPAR cell line from a single-cell clone of a KP tumour expressing APOBEC3B which had developed in an immune-deficient background and therefore could not undergo immune-editing. We used the KPAR cell line as an orthotopic transplantable model of lung cancer and demonstrated that this model was immunogenic, stimulating anti-tumour immune responses which sensitised tumours to immune checkpoint blockade. Although the cell line was generated from the KP-A3Bi GEMM, it did not possess a substantial number of new mutations and instead we observed anti-tumour immune responses directed against derepressed endogenous retroviral proteins. Several immunogenic murine cancer cell lines, including CT26^35^, B16 and MC38^36^, have been shown to express endogenous retroviral proteins which can act as antigens that are recognised by T cells. Similarly, anti-tumour immune responses have been observed against human endogenous retroviruses in melanoma^37^ and breast cancer^37^, however the relevance of these antigens in lung cancer remains unclear. Nevertheless, elevated expression of a tumour-associated antigen may better model anti-tumour immune responses that occur in human cancers compared to the expression of strong foreign antigens such as ovalbumin, luciferase^9^ or lymphocytic choriomeningitis virus glycoprotein^38^. Indeed, the expression of these antigens in KP lung cancer cells results in the rejection of transplanted cells or the selection of clones that have lost antigen expression^9^.

Most studies of immunotherapy utilising transplantable cell lines involve subcutaneous transplantation into syngeneic immune-competent mice. We observed a striking difference in the response to anti-PD-1 or anti-CTLA-4 in subcutaneous versus orthotopic tumours, as previously demonstrated for PD-1/PD-L1 blockade^26^. Subcutaneous KPAR tumours responded to anti-CTLA4 but were refractory to anti-PD-1, as observed in an immunogenic melanoma model^39^. Anti-CTLA-4 can induce tumour regression through the depletion of Tregs in subcutaneous tumours^25^. We validated this observation in subcutaneous KPAR tumours but observed that anti-CTLA-4 was not sufficient to deplete Tregs in the orthotopic setting. Furthermore, the ratio of Foxp3^+^ Tregs to CD4^+^ effector T cells was much higher in the subcutaneous setting. This may explain why subcutaneous tumours are refractory to PD-1 blockade but respond to anti-CTLA-4. Conversely, the reduced fraction of Tregs in orthotopic tumours may explain the minimal response to CTLA-4 blockade. Together these results highlight the importance of studying tumours in their tissue of origin when assessing responses to ICB, with orthotopic tumours likely to yield more directly clinically relevant information than subcutaneous tumours.

The recently developed KRAS^G12C^ specific inhibitors have produced outstanding responses in NSCLC^40^. A recent study showed that these inhibitors could promote T cell responses through increased IFNγ signalling in the immunogenic CT26^G12C^ colon cancer transplantable model^7,41^. Similar to what was shown in these studies, we observed profound changes in the tumour microenvironment in response to KRAS^G12C^ inhibition, indicative of enhanced anti-tumour immune responses. Furthermore, the tumour regression we observed after KRAS^G12C^ inhibition in KPAR^G12C^ tumours was more profound in immune-competent mice; however, this was not the case for non-immunogenic KPB6^G12C^ tumours. This result suggests that the efficacy of KRAS^G12C^ inhibitors is partially due to the engagement of the adaptive immune system in immune-hot tumours. The KPAR model therefore offers the possibility to explore combinations of immunotherapy with KRAS^G12C^ inhibition to overcome the acquired resistance anticipated following this novel targeted therapy^42^.

In conclusion, we have created a novel model of immunogenic KRAS-driven lung adenocarcinoma, which we anticipate will contribute to the development of new combinations of therapies, including those involving immune checkpoint blockade and KRAS^G12C^ inhibition.

## METHODS

### In vivo tumour studies

*Kras*^LSL-G12D/+^;*Trp53*^fl/fl^ mice (KP) were sourced from the Mouse Models of Human Cancer Consortium and maintained on a pure C57Bl/6J background. *Kras*^LSL-G12D/+^; *Trp53*^fl/fl^;*Rosa26*^*A3Bi*^ mice (KPA) and *Kras*^LSL-G12D/+^;*Trp53*^fl/fl^;*Rosa26*^*A3Bi*^;*Rag1*^KO/KO^ mice (KPAR) were generated by breeding KP mice with *Rosa26*^*A3Bi*^ mice and *Rag1*^KO/KO^ mice (see supplement for development and validation of the *Rosa26::LSL-A3Bi* model). Tumours were induced by intratracheal intubation of 1 × 10^6^ adenovirus expressing Cre recombinase as previously described^43^. Tumour volume was assessed via micro-CT scanning.

For the urethane-induced models, tumours were induced by 3 intra-peritoneal injections of 1 mg/g of urethane over the period of a week. Three weeks following urethane first injection, APOBEC3Bi was induced by 3 doses of 100 mg/g tamoxifen over a period of a week in *Rosa26*^*A3Bi/CreER(t2)*^ mice (UrA3Bi-CreER). Tumour volume was assessed via micro-CT scanning.

All transplantation animal experiments were carried out using 8-12-week C57BL/6J mice. For subcutaneous studies, 1.5 × 10^5^ KPAR1.3 or KPAR1.3^G12C^ cells and 5 × 10^5^ KPB6^G12C^ cells (1:1 mix with matrigel) were injected subcutaneously into the flank. Tumour volume was measured twice weekly and calculated using the formula 0.5 × [Length × Width^2^]. Mice were euthanised when the average tumour dimensions exceeded 1.5 mm. For re-challenge experiments, mice that remained tumour-free for 4 months were injected subcutaneously into the opposite flank with 1.5 × 10^5^ KPAR1.3 tumour cells. For orthotopic studies, 1.5 × 10^5^ KPAR1.3 cells, 1 × 10^5^ KPAR1.3^G12C^ and KPB6 cells were injected intravenously into the tail-vein. Mice were euthanised when the humane endpoint of 15% weight loss was reached.

For treatments, 200 μg anti-PD-1 (clone RMP1-14, BioXcell) and 200 μg anti-CTLA-4 (clone 9H10, BioXcell), or their respective IgG controls, were administered per mouse via intraperitoneal injection twice weekly for a maximum of three weeks. Anti-PD-L1 (clone 10F.9G2, BioXcell) or the respective IgG control were administered at 10 mg/kg via intraperitoneal injection twice weekly, for two weeks. AZ-8037 or vehicle (10% Pluronic-F127) was administered 5 days per week via oral gavage at 100 mg/kg. Mice were randomised into groups and treatments initiated once tumours reached an average volume of 150 mm^3^ for subcutaneous studies or were detectable by micro-CT for orthotopic experiments.

### Cell lines

The KPB6 cell line was obtained from Cell Services at the Francis Crick Institute. KPAR and KPA cell lines were established by cutting up lung tumours into small pieces and culturing in DMEM-F12 supplemented with Glutamax^®^, FBS (10%), hydrocortisone (1 μM), EGF (20 ng/ml), IGF (50 ng/ml), penicillin (100 units/mL) and streptomycin (100 μg/mL). KPAR, KPA and KPB6 cell lines were cultured in DMEM supplemented with fetal bovine serum (10%), L-glutamine (2 mM), penicillin (100 units/mL) and streptomycin (100 μg/mL). Clonal cells were derived by single-cell dilution into 96 well plates. Cell lines were routinely tested for *Mycoplasma*.

### Flow cytometry

Mouse tumours were cut into small pieces, incubated with collagenase (1 mg/ml; ThermoFisher) and DNase I (50 U/ml; Life Technologies) for 45 min at 37°C and filtered through 70 μm strainers (Falcon). Red blood cells were lysed for 5 min using ACK buffer (Life Technologies). Cells were stained with fixable viability dye eFluor870 (BD Horizon) for 30 min and blocked with CD16/32 antibody (Biolegend) for 10 min. Cells were then stained with one of three antibody cocktails for 30 min (see Supplementary Table 1). Intracellular staining was performed using the Fixation/Permeabilization kit (eBioscience) according to the manufacturer’s instructions. Samples were resuspended in FACS buffer and analysed using a BD Symphony flow cytometer. Data was analysed using FlowJo (Tree Star).

For FACS analysis *in vitro*, cells were trypsinised, washed with FACS buffer and stained for eMLV envelope glycoprotein using the 83A25 monoclonal antibody followed by a secondary staining with anti-rat IgG2a (PE). Samples were run on LSRFortessa (BD).

| Mix 1 | Mix 2 | Mix 3 |
| --- | --- | --- |
| CD3 [17A2], FITC | CD45 [30-F11], PerCP | MHC-II [M5/114.15.2], FITC |
| CD45 [30-F11], PerCP | CD49b [DX5], AF488 | CD45 [30-F11], PerCP |
| Foxp3 [FJK-16s], eFluor660 | NKp46 [29A1.4], BV421 | TIM3 [RMT3-23], PE |
| CD44 [IM7], BV421 | F4/80 [EMR1], BV785 | CD103 [M290], BV421 |
| CD69 [JES5-16E3], BV605 | Ly6G [1A8], BV711 | CD24 [M1/69], BV605 |
| CD62L [MEL-14], BV711 | Ly6C [HK.14], BV785 | CD206 [C068C2], BV711 |
| PD-1 [29F.1A12], BV785 | CD11c [HL3], BUV395 | CD86 [GL-1], BV785 |
| CD8 [53-6.7], BUV395 | B220 [RA3-6B2], BV605 | CD11c [HL3], BUV395 |
| CD4 [GK1.5], BUV737 | CD11b [M1/70], BUV395 | CD11b [M1/70], BUV737 |
| TIM-3 [RMT3-23], PE | CD19 [6D5], PE | PD-L1 [MIH5], PE |
| LAG-3 [eBioC9B7W], PE-Cy7 | PD-L1 [MIH5], PE-Cy7 | CD64 [X54-5/7.1], PE-Cy7 |

### Histopathology and Immunohistochemistry

Tumour-bearing lungs were fixed in 10% NBF for 24 h followed by 70% ethanol. Fixed tissue was embedded in paraffin wax. Tissue sections were stained with haematoxylin and eosin, using standard methods. Sections were examined by two board-certified veterinary pathologists (ASB and SLP). The tumour grade (adenoma, well-, or moderately well-differentiated carcinoma) was defined as follows, adenoma; mild pleomorphism, anisocytosis and anisokaryosis and well-circumscribed, well-differentiated carcinoma; mild to moderate pleomorphism, anisocytosis and anisokaryosis and increased mitotic activity, and moderately well-differentiated carcinoma; as well-differentiated carcinoma but with evidence of invasion.

For immunohistochemistry staining, tissue sections were boiled in sodium citrate buffer (pH 6.0) for 15 min and incubated with the following antibodies for 1h: anti-Foxp3 (D6O8R, CST), anti-CD8 (4SM15, Thermo Scientific) and anti-A3Bi (5210-87-13)^44^. Primary antibodies were detected using biotinylated secondary antibodies and detected by HRP/DAB. Slides were imaged using a Leica Zeiss AxioScan.Z1 slide scanner.

### Micro-CT imaging

Mice were anesthetised by inhalation of isoflurane and scanned using the Quantum GX2 micro-CT imaging system (Perkin Elmer) at a 50μm isotropic pixel size. Serial lung images were reconstructed and tumour volumes subsequently analysed using Analyse (AnalyzeDirect).

### Exome sequencing and neoantigen prediction

Genomic DNA was extracted from tumours, cell lines and mouse tails using DNeasy Blood & Tissue kit (Qiagen). Exome libraries were prepared by the Advanced Sequencing Facility at the Crick and sequenced using an Illumina HiSeq 4000 using 100 base pair paired-end reads. We aligned the reads to the GRCm38 mouse genome (Ensembl, release-86) using bwa mem (v0.7.15)^45^. We processed all alignment files using samtools (v1.4)^46^. We identified and marked duplicates using Picard MarkDuplicates (v2.1.1, Java v1.8.0_92). We realigned indel regions using GATK RTCreator and indelRealign (v4.1.3.0, Java v1.8)^47^. We called germline mutations in each of the normal tail samples using Mutect2 (GATK v4.1.3.0). We used these tail germline calls to construct a ‘panel of normals’ (PON) which we used for downstream somatic mutation calling. We called tumour specific somatic mutations against matched tail samples from the same animal using Mutect1 (v1.1.7) and Mutect2 (GATK v4.1.3.0). We used C57BL mouse strain germline SNPs from the Mouse Genome Project^47^ to filter for known germline variants. (Mutect1 –dbsnp; Mutect2 SelectVariants –discordance, gatk CreateSomaticPanelOfNormal --germline-resource). We created a PON using gatk GenomicsDBImport and CreateSomaticPanelOfNormal (v4.1.3.0, Java v1.8). We generated Mutect2 calls using gatk Mutect2, FilterMutectCalls and SelectVariants sub-commands. We annotated SNVs using the predictCoding() function from the VariantAnnotation Bioconductor package (v1.26.1, R v3.5.1). We identified regions of relative copy number variation between the tumour and matched tail samples using cnvkit batch^48^ (v0.9.5, default parameters) (annotation as above).

We corrected variant allele frequencies for tumour purity using two different methods depending on the model. In the case of the KP induced tumours we calculated the ratio of reads per base across the floxed exons of Trp53 (1-9) and the undeleted exon 11. In the cases of the Urethane induced models we used the VAF of the Kras Q61 mutation. We assumed this mutation to be the primary driver of tumour activation since it is found in all the urethane induced tumours. All Kras Q61 loci were found to be copy number neutral. We called absolute copy number counts from the relative ratios using cnvkit call. The ratios were scaled using the purity estimates calculated above and integer copy numbers assigned using the follow log2 ratio thresholds, with used these tumour purity estimates and the relative copy number ratios to estimate the clonality of SNVs. In the case of KPA(R) cell lines, we assumed clonality of most CNAs and scale the CN to the X chromosome (haploid in male cell lines and diploid in female cell lines).

We distinguished clonal from sub-clonal mutations by estimating the fraction of tumour cells carrying the observed mutation using the method detailed in Turajlic *et al.*^49^. Briefly we calculated the expected VAFs, given a tumour purity estimate, across a range of tumour cell fractions carrying the mutation [00.1, 00.2 ... 1] and possible mutation copy numbers ranging from 1 to the reported absolute copy number reported above. We selected the tumour cell fraction and mutation copy number that gave an expected VAF closest to the observed. In situations where the expected VAFs for different mutation copy numbers were all close to the observed the more likely mutant copy number was selected. We classified an SNV as clonal if it was found in > 75% of the tumour cells, subclonal if less. We corrected purity normalized VAFs for CNV so reported VAFs were assumed to be diploid.

Mutated peptide sequences were processed using NetMHC4.0 with *k*-mer of 8-11 length. Rank threshold of 0.5, or 2.0, were used to identify putative strong, or weak, MHC class I (H2-Kb and H2-Db) binders, respectively.

### qPCR

Total RNA was extracted from cell lines or frozen tumour samples using RNeasy kit (Qiagen). Frozen tumour samples were homogenised prior to RNA extraction either using a syringe and needle or QIAshredder columns (Qiagen). cDNA was synthesised using Maxima First Strand cDNA Synthesis Kit (Thermo Fisher Scientific) and qPCR was performed using Applied Biosystems™ Fast SYBR™ Green Master Mix (Thermo Fisher Scientific). mRNA relative quantity was calculated as previously described^50^ and normalised to at least three housekeeping genes.

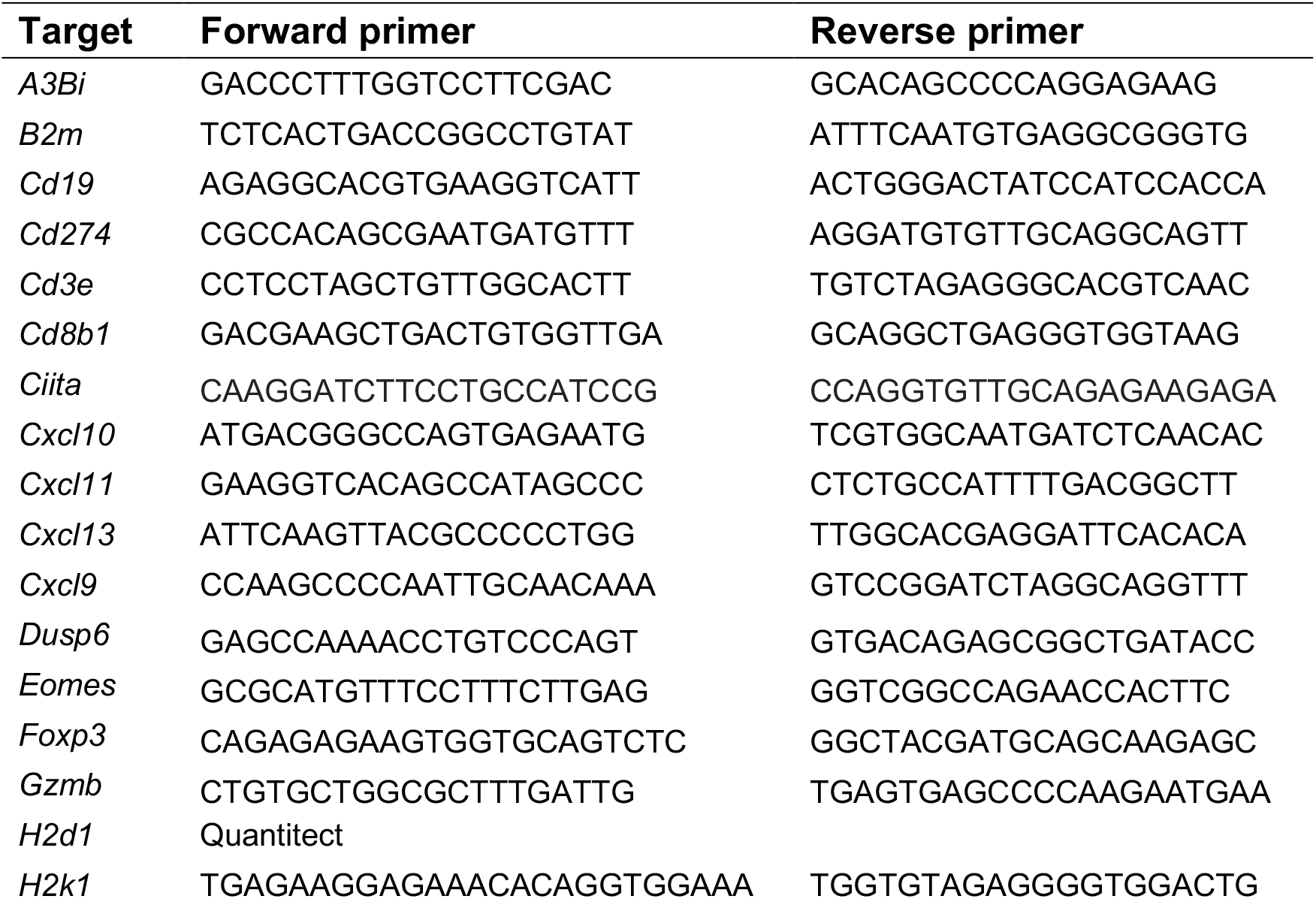
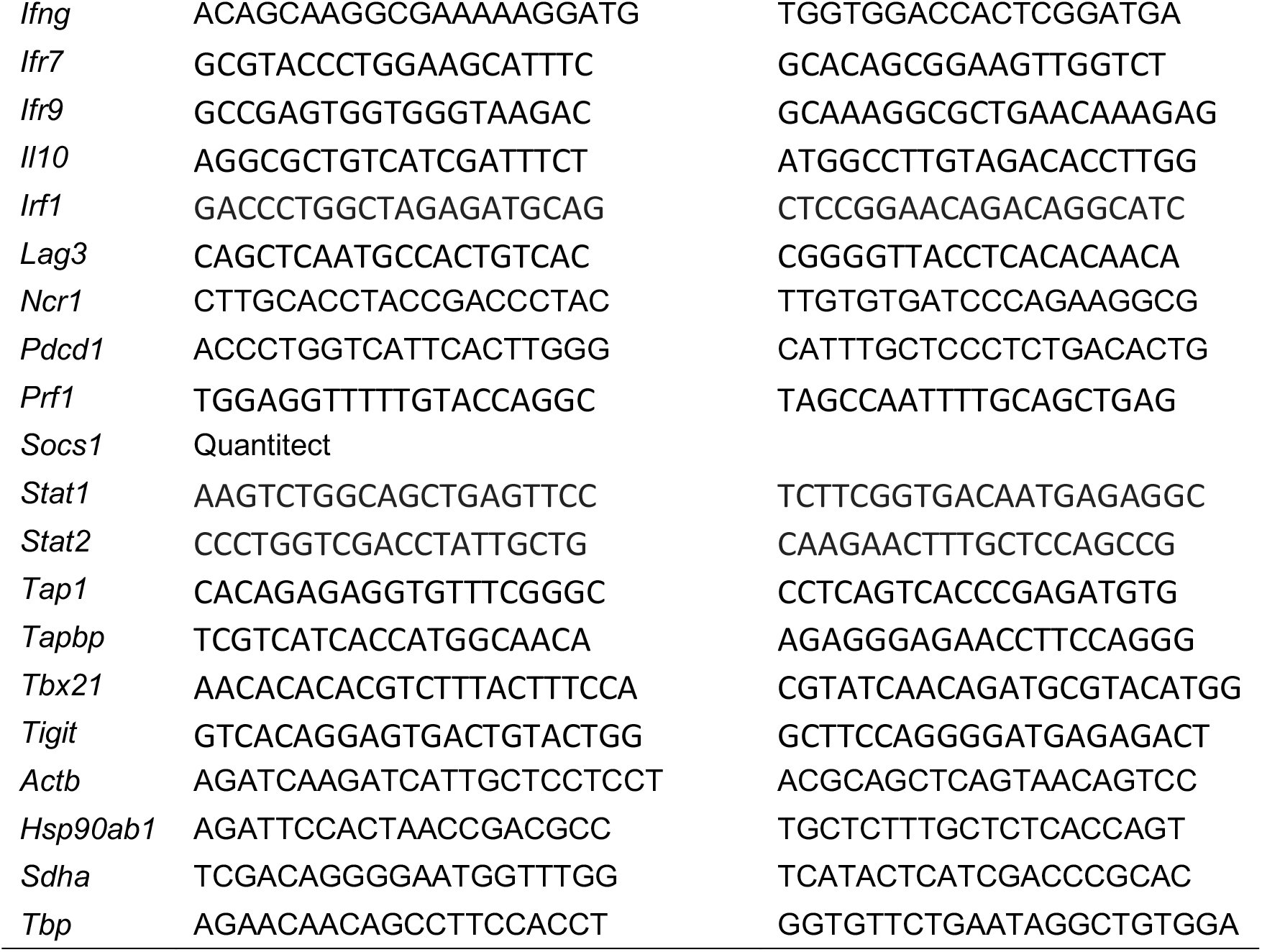

### Tumour Cell Viability

For short-term viability assays, 1.5 × 10^3^ KPAR1.3^G12C^ or 2 × 10^3^ KPB6^G12C^ cells were seeded in 96-well plates and grown in the presence of different inhibitors for 72 h. Cell viability was assessed using CellTiter-Blue (Promega).

### Western blotting

Cells were lysed using protein lysis buffer (Sigma) with protease and phosphatase inhibitor cocktails (Sigma). Protein concentration was determined using a BCA protein assay kit (Pierce). 15-20 μg of protein was separated on a 4-12% Bis-Tris gel (Life Technologies) and transferred to PVDF membranes. Protein expression was detected by Western blotting using the following primary antibodies against: S6 (54D2, Cell Signalling), p-S6 (Ser235/236) (2211, Cell Signalling), Erk1/2 (3A7, Cell Signalling), p-Erk1/2 (Thr202/Tyr204) (9101, Cell Signalling), Akt (40D4, Cell Signalling), p-Akt (Ser473) (D9E, Cell Signalling), and Vinculin (VIN-11-5, Sigma). Primary antibodies were detected with HRP-conjugated anti-rabbit or anti-mouse IgG and visualised with Immobilon Western HRP substrate (Merck).

### ELISpot analysis

1 × 10^4^ CD8+ TILs, isolated from tumours using the EasySep Mouse CD8α Positive Selection Kit (Stemcell Technologies), were harvested from tumour-bearing mice and pulsed with 1 μM peptide corresponding to neoantigens predicted from KPAR1.3 WES (Supplementary Table 1) or eMLV *env* peptide (KSPWFTTL). TILs were co-incubated with 1 × 10^5^ splenocytes from naïve mice as a source of dendritic cells. Cells were stimulated for 24h in anti-mouse IFNγ-coated ELISpot plates (BD Bioscience). Plates were developed according to manufacturer’s instructions and quantified using a CTL S6 machine.

## ACKNOWLEDGEMENTS

We thank the science technology platforms at the Francis Crick Institute including Biological Resources, Scientific Computing, Bioinformatics and Biostatistics, Flow Cytometry, Experimental Histopathology, and Cell Services. We also thank Colleen Forster and Gerard O’Sullivan for assistance with immunohistochemistry and pathology, Yasuhiko Kawakami for sharing CMV-Cre animals, and Brian Dunnette for expertise with the Aperio ScanScope XT at the University of Minnesota.

## Funding

This work was supported by funding to J.D. from the Francis Crick Institute— which receives its core funding from Cancer Research UK (FC001070), the UK Medical Research Council (FC001070), and the Wellcome Trust (FC001070)—from the European Research Council Advanced Grant RASImmune, and from a Wellcome Trust Senior Investigator Award 103799/Z/14/Z. The *A3Bi* minigene model was developed with support from the National Cancer Institute P01-CA234228 (RSH), Team Judy (RSH), Randy Shaver Cancer Research and Community Fund (RSH), University of Minnesota Masonic Cancer Center, and College of Biological Sciences (RSH). RSH is the Margaret Harvey Schering Land Grant Chair for Cancer Research, a Distinguished University McKnight Professor, and an Investigator of the Howard Hughes Medical Institute. S.C.T received funding from the European Union’s Horizon 2020 research and innovation programme under the Marie Sklodowska-Curie grant agreement No 703228

## Competing interests

J.D. has acted as a consultant for AstraZeneca, Bayer, Jubilant, Theras, Vividion and Novartis. R.S.H is a co-founder, shareholder, and consultant of ApoGen Biotechnologies Inc. S.R. is an employee of AstraZeneca. C.S. receives grant support from Archer Dx, AstraZeneca, Boehringer–Ingelheim and Ono Pharmaceutical; has consulted for AstraZeneca, Bicycle Therapeutics, Celgene, Genentech, GRAIL, GSK, Illumina, Medicxi, MSD, Novartis and the Sarah Cannon Research Institute; receives grant support and has consulted for Bristol Myers Squibb, Pfizer and Roche–Ventana; is an advisory board member and is involved in trials sponsored by AstraZeneca; has stock options in Apogen Biotechnologies, Epic Sciences, GRAIL; and has stock options and is a co-founder of Achilles Therapeutics. The other authors declare that they have no competing interests.

## Author contributions

S.C.T, J.B., E.K.L., G.K., R.S.H. and J.D. designed the study, interpreted the results and wrote the manuscript. S.C.T, J.B., P.R.C., M.M-A., M.A.C., S.R., E.M., K.N. and D.H. performed the biochemical experiments, C.M. assisted with *in vivo* studies, P.P.A., C.D., A.S.B., S.P. performed pathological studies, S.H. and P.E. performed bioinformatics analyses, R.S.H. designed the A3B knock-in model, supervised validation studies, contributed support and reagents, obtained funding, and edited the manuscript. E.K.L. created DNA constructs, performed validation studies, and conducted full body expression experiments. W.L.B., L.K.L., and C.D. provided technical and logistical assistance. R.I.V. provided statistical analysis. All authors contributed to manuscript revision and review.

## Notes

### Summary of Updates

This version of the manuscript has been revised to reflect our finding that endogenous CD8+ T cell responses are directed against derepressed endogenous retrovirus antigens expressed by the KPAR1.3 tumour cell line, but evidence of T cell responses against APOBEC3B induced mutant neoantigens was not found.

