## Supplemental Figures 1-8, Supplemental methods for "An immunogenic model of KRAS-mutant lung cancer for study of targeted therapy and immunotherapy combinations"

### SUPPLEMENTARY INFORMATION

### Boumelha et al.

#### Supplementary Methods

##### *Rosa26::LSL-A3Bi* knock-in construction

An intron-containing human *A3B* cDNA (*A3Bi*)[1] was amplified using 5'-**ACG-CGT-**ACC-ATG-AAT-CCA-CAG-A and 5'- **ACG-CGT-**tca-GTT-TCC-CTG-ATT-C (*Mlu*I sites in bold) and cloned into pJET2.1 (Clontech). The resulting cDNA was subcloned into a *Mlu*I-digested *Rosa26* gene targeting vector pROSA26-1 (gift from Philippe Soriano; Addgene plasmid #21714; http://n2t.net/addgene:21714; RRID:Addgene_21714)[2] which placed *A3Bi* downstream of a *loxP*-flanked transcription stop cassette (schematic in **Supplementary Fig. 2A**). The functionality of this construct was confirmed by directional subcloning into the *Nhe*I/*Cla*I sites of pcDNA3.1 (ThermoFisher Scientific) followed by co-transfecting 293T cells with a control vector or Cre recombinase expression plasmid (TransitLT-1, Mirus) and, 48 hrs later, performing *A3B*-specific RT-qPCR, anti-A3B immunoblots, and single-stranded DNA C-to-U activity assays (procedures described below; representative data in **Supplementary Fig. 2B, C**). Additional experiments showed that the hallmark nuclear localization of human A3B in human cells[3–5] is also evident upon expression in murine NIH 3T3 cells (**Supplementary Fig. 2D**). 10 µg of functionally confirmed targeting vector was linearized with *Sac*II and electroporated into C57BL/6N embryonic stem (ES) cells (Ingenious Targeting Laboratory, Ronkonkoma, NY). G418 was used to select clones and PCR screening was used to identify homologous recombinants (upstream forward primer RSH8979 5'-ACT-GAC-CGC-ACG-GGG-ATT-CC and *neo* reverse primer RSH8985 5'-ACT-GAC-CGC-ACG-GGG-ATT-CC). Positive ES clones were confirmed by digesting genomic DNA with *Mfe*I and *Bsr*G1, separating on a 0.8% agarose gel, transferring to nylon membrane, and Southern blotting (restriction sites and probe locations shown in **Supplementary Fig. 2A** and blot images in **Supplementary Fig. 2E, F**). Correctly targeted ES clone 243 was microinjected into BALB/c blastocysts, and chimeric pups with high percentages of black coat colour were mated to wildtype C57BL/6J mice to obtain germline transmission.

**Animal husbandry and genotyping**

Mice were maintained at the University of Minnesota in AAALAC-accredited animal facilities (IACUC protocol 1302A30327). Heterozygous *Rosa26::LSL-A3Bi* (2x *loxP*) female mice were crossed with CMV-Cre male animals to remove the transcription stop cassette and obtain *Rosa26::L-A3Bi* (1x *loxP*) progeny (Jackson Lab Stock No: 006054 provided by Yasuhiko Kawakami, University of Minnesota). Cre-mediated recombination was verified by a PCR-based strategy (representative data in **Supplementary Fig. 2G**; methods detailed below). Because the CMV-Cre transgene is located on the X-chromosome, *Rosa26::L-A3Bi* (1x *loxP*) males were bred with wildtype females to generate a cohort with whole-body A3B expression. Human A3B expression was demonstrated in multiple tissues by RT-qPCR and single-strand DNA deaminase activity assays (**Supplementary Fig. 2H**). Protein-level expression and nuclear localization was also demonstrated by immunofluorescent microscopy of A3B in MEFs and anti-A3B IHC of a wide range of tissues including lung (procedures described below and representative data in **Supplementary Fig. 2I, J**. Expression of human A3B from the *Rosa26* promoter did not compromise fertility, as evidenced by normal Mendelian progeny ratios (not shown). Moreover, whole-body expression of A3B caused no overt somatic phenotypes in otherwise wildtype C57BL/6J animals. Mice were monitored weekly for tumour formation and euthanized by $CO_2$ asphyxiation upon natural health decline. The overall life expectancy of animals with whole-body expression of A3B was indistinguishable from animals without Cre-mediated induction of the minigene (overall KM plots in **Supplementary Fig. 2K**). Moreover, rates of tumour occurrence were similar between the two groups (KM plots of tumour-free survival in **Supplementary Fig. 2L**). Tumours were rare and average tumour numbers were similarly low in A3B-expressing versus non-expressing controls (dot plots in **Supplementary Fig. 2M**). Genomic DNA was isolated from tail biopsies of 21-day-old mice using the Gentra Puregene protocol (Qiagen) and 50 ng was used as a template for each diagnostic PCR reaction. The *Rosa26::LSL-A3Bi* (unrecombined 2x *loxP* allele) was detected by PCR of a 500 bp fragment using primers RSH8980 5’-AGC-ACT-TGC-TCT-CCC-AAA-GTC (*Rosa26* forward) and RSH8985 and 5’-TGC-GAG-GCC-AGA-GGC-CAC-TTG-TGT-AGC (LSL cassette reverse). The *Rosa26::L-A3Bi* (recombined 1x *loxP* allele) was detected by PCR of a 598 bp fragment using the same *Rosa26* forward primer and RSH8984 5'-GCA-CAT-TTC-TGC-GTG-GTA-CTG-AGG (A3B reverse). The wildtype *Rosa26* locus was detected as a 300 bp PCR product using the same *Rosa26* forward primer and RSH10347 5'-CAC-CTG-TTC-AAT-TCC-CCT-GC-3' (*Rosa26* reverse). CMV-Cre was detected by standard PCR of a 100 bp fragment using primers 5'-GCG-GTC-TGG-CAG-TAA-AAA-CTA-TC (RSH7141) and 5'-GTG-AAA-CAG-CAT-TGC-TGT-CAC-TT (RSH7142). A representative agarose gel image is shown in **Supplementary Fig. 2g**.

**Cell culture**

293T, NIH 3T3, and primary MEFs were cultured at 37°C containing 5% $CO_2$ and maintained in Dulbecco's Modified Eagle Media (DMEM) supplemented with 10% fetal bovine serum (FBS), 100 U/mL penicillin, 100 µg/mL streptomycin. ES cells were plated on irradiated primary MEF feeder cells plated on 0.1% gelatin coated plates. ES cells were cultured at 37°C containing 7.5% $CO_2$ and fed daily with IMDM supplemented with 15% FBS, 1x sodium pyruvate (Invitrogen), 1x L-glutamine (Invitrogen), 1x non-essential amino acids (Sigma-Aldrich), 100 U/mL penicillin, 100 µg/mL streptomycin, 10 ng/ml LIF (Millipore), and 1x 2-mercaptoethanol (ThermoFisher Scientific).

**RT-qPCR for quantification of *A3B* mRNA levels**

Total RNA was isolated with the Qiagen RNAeasy protocol with QIAshredder and on column DNase treatment. cDNA was prepared and *A3B* mRNA expression was quantified by RT-qPCR as described[5]. Briefly, reverse transcription of 1 µg total RNA was done using Transcriptor Reverse Transcriptase (Roche) and the resulting cDNA was diluted 1:5 and used as template for quantitative PCR. *A3B* expression was quantified using a LightCycler 480 (Roche) with forward primer 5'-GAC-CCT-TTG-GTC-CTT-CGA-C (RSH3220) and reverse primer 5'-GCA-CAG-CCC-CAG-GAG-AAG (RSH3221) and detected using Roche UPL #01. The housekeeping gene, human *TBP* or murine *Tbp*, was used for gene expression normalization between samples. Human *TBP* was amplified using primers 5'-CCC-ATG-ACT-CCC-ATG-ACC (RSH3231) and 5'-TTT-ACA-ACC-AAG-ATT-CAC-TGT-GG (RSH3232) and detected using Roche UPL #51. Murine *Tbp* was amplified using primers 5'- GGG-GAG-CTG-

TGA-TGT-GAA-GT (RSH2913) and 5'- CCA-GGA-AAT-AAT-TCT-GGC-TCA (RSH2914) and detected using Roche UPL #97.

**A3B immunoblots and single-stranded DNA C-to-U activity assays**

Anti-A3B immunoblots were performed using established procedures[6–9]. Whole cell extracts were prepared by boiling cell suspensions for 30 min in 2.5x reducing sample buffer (31 mM Tris pH 6.8, 10% glycerol, 1% SDS, 1.25% 2-mercaptoethanol, and 0.05% bromophenol blue). Soluble proteins from 30,000 cells were fractionated by SDS-PAGE (12.5% polyacrylamide gel), transferred to PVDF membrane, and probed with the rabbit anti-A3B mAb 5210-87-13 (1:1000; Harris lab custom reagent)[6]. A murine anti-tubulin mAb was used as a loading control (1:40,000; Novus Biologicals). Licor secondary antibodies, anti-rabbit IRdye 800CW (Licor 827-08365) and anti-mouse IRdye 680LT (Licor 925-68020), were used at 1:20,000 and visualized using a Licor Odyssey imaging system.

Cell extracts were prepared for single-stranded DNA C-to-U activity assays as described[7–10]. Cells were suspended in HED buffer (25 mM HEPES, 5 mM EDTA, 10% glycerol, 1 mM DTT, 1 tablet proteasome inhibitor (Roche) per 50 ml buffer), freeze/thawed once, and vortexed to promote lysis. Cell debris was removed by centrifugation and cleared lysates were incubated 2 hrs with 4 pmol of a fluorescently labelled ssDNA substrate with a single target cytosine (5'-ATT-ATT-ATT-ATT-CGA-ATG-GAT-TTA-TTT-ATT-TAT-TTA-TTT-ATT-T-fluorescein; RSH5195). These conditions support A3B dependent ssDNA cytosine deamination and excision of the resulting uracil by uracil DNA glycosylase. Cleavage of uracil-excised substrates was promoted by adding NaOH to a final concentration of 0.1M and incubating samples at 98°C for 5 min. The resulting samples were fractionated on a 15% acrylamide gel, imaged (Typhoon, GE Healthcare Life Sciences), and quantified by densitometry (ImageQuant).

**A3B immunohistochemistry (IHC)**

IHC staining was performed following described procedures[6,11,12]. FFPE tissues were sectioned at 4 µm, mounted on positively charged, adhesive slides and allowed to air-dry for at least 24 hrs. To deparaffinize and rehydrate the samples, slides were baked in a 65°C oven for 20 min, washed 3 times with CitrisolvTM (Decon Labs, #1601) for 5-min/each, soaked in graded alcohols (100% x 2, 95% and 80% for 3 min/each), and then rinsed in running water for at least 5 min. Epitope retrieval was performed using Reveal Decloaker (BioCare Medical, #RV1000M) in a steamer for 35 min, followed by a 20 min “cool-down” period. Then, slides were rinsed with running tap water for 5 min and transferred to TBST for 5 min. Endogenous peroxidase activity was quenched by placing the slides in 3% H2O2 in TBST for 10 min at RT, followed by a 5-min rinse under running water. To block non-specific binding of primary antibody, sections were covered with Background Sniper (BioCare Medical, #BS966MM) for 15 min at RT. After blocking, serial sections of each specimen were incubated overnight at 4°C with a rabbit anti-human A3B mAb (5210-87-13)[6] diluted 1:350 in 10% Sniper in TBST. This mAb does not cross-react with murine A3.

Following overnight incubation with primary antibody, sections were rinsed in TBST for 5 min, and completely covered with anti-rabbit poly-HRP-IgG (Leica Biosystems, Novolink Polymer, #RE7260-K) for 30 min at RT. The reaction product was developed using the Novolink DAB substrate kit (Leica Biosystems, # RE7230-K) at RT for 3-5 min, rinsed in tap water for 5 min, counterstained in Mayer’s hematoxylin solution (Electron Microscopy Sciences, #26252-01) for up to 5 min, dehydrated in graded alcohols and CitrisolvTM, and cover-slipped using Permount mounting media. The stained slides were scanned at 40x magnification and A3B nuclear immunoreactivity was visualized with the Aperio ScanScope XT (Leica Biosystems).

**Gene editing**

Double-stranded DNA breaks were induced at the *Kras* locus in the KPB6 cell line by nucleofection (Amaxa Nucleofector kit V, protocol T-013) of the pX458 vector, as previously described[13] using the following sgRNA: 5’-CTTGTGGTGGTTGGAGCT GA-3’. Homologous recombination repair was induced using the following single-stranded DNA template: 5’-ATTTAGTTGTATTTTATTATTTTTATTGTAAGGCCTG CTGAAAATGACTGAGTATAAGCTAGTCGTATTGGAG CTTGTGGCGTAGGCAA GAGCGCCTTGACGATACAGCTAATTCAGAATCACTTTGTGGATGAA-3’0.

Prime editing technology was used to induce single-strand DNA break and reverse transcriptase-mediated genome editing in KPAR1.3 cells, as previously described[14]. Golden Gate Assembly was used to clone the sgRNA (5’TCAGCTCCA ACCACCACAAG-3’), 3’ extension sequence: (GCCTACGCCACAAGCTCCAACC ACCA-3’) and sgRNA scaffold (5’-CTAGAAATAGCAAGTTAAAATAAGGCTAG TCCGTTATCAACTTGAAAAAGTGGCACCGAGTCG-3’) into a mammalian U6 expression vector (Addgene, #132777). To increase the efficiency of editing, a second sgRNA lacking the 3' extension sequence and targeting upstream of the editing point was generated by cloning the following sgRNA: (5'-TATACTCAGTCAT TTTCAGC-3') and identical sgRNA scaffold sequence into the U6 vector. Guide-expressing plasmids and a Cas9-H840A/GFP-expressing plasmid (Addgene, #132776) were transfected into KPAR1.3 cells by nucleofection (Amaxa Basic Nucleofector Kit for Primary Mammalian Epithelial Cells, protocol T-030). All DNA oligos were ordered annealed and phosphorylated from Integrated DNA Technologies and overhang sequences were included to make them suitable for Golden Gate assembly.

For both KPAR1.3 and KPB6 cells, genomic DNA from single cell clones was extracted using QuickExtract DNA Extraction Solution (Lucigen) and amplified with PrimeSTAR Max DNA polymerase (Takara Bio) using primers Kras-F: 5'-GTCC ACAGGGTATAGCGTACT-3' and Kras-R 5'-CACCCAGTTTAAAGCCTTGGA-3'. PCR products were digested for 1h at 37°C with BfaI (NEB) for the KPB6 cell line or BccI (NEB) for KPAR1.3 cells. Digestion products were evaluated by gel electrophoresis with a 2% agarose gel.

Genotypes of single-cell clones were verified by Sanger sequencing using the following primer: 5'-CACCCAGTTTAAAGCCTTGGA-3'. Alternatively, Illumina MiSeq next-generation sequencing was used. For this purpose, genomic DNA was amplified with CloneAmp HiFi PCR Premix (Takara Bio) using primers Kras1-F: 5'-TCGTCGGCAGCGTCAGATGTGTATAAGAGACAGGTCCACAGGGTATAGCGT ACT-3' and Kras1-R: 5'-GTCTCGTGGGCTCGGAGATGTGTATAAGAGACAGTTAC

AAGCGCACGCAGACT-3'. PCR products were purified using AMPure XP beads according to the manufacturer's instructions and sequenced on the Illumina MiSeq. All sequences were visualized and analysed using SnapGene Software.

#### Supplementary figure 1

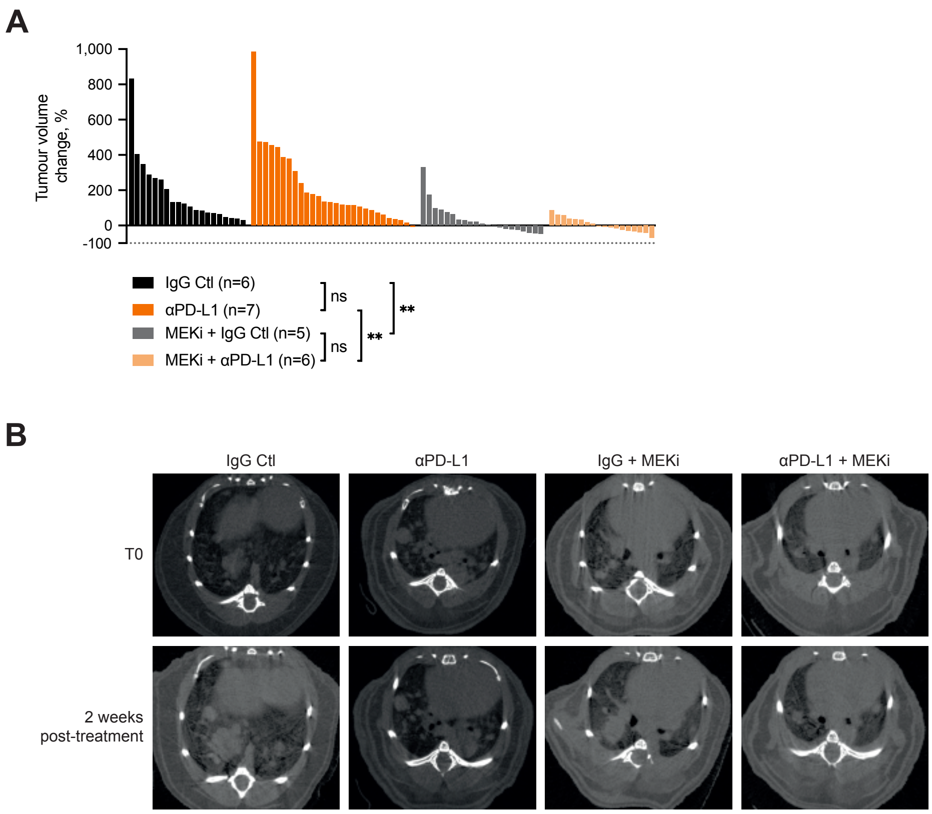


**Supplementary figure 1. MEK inhibition does not improve response to anti-PD-L1**
(A) Waterfall plot of tumour volume change in KP-tumour-bearing mice treated intraperitoneally with anti-PD-L1 (10 mg/kg) or corresponding isotype control (IgG Ctl) twice weekly for 2 weeks and/or trametinib (3 mg/kg daily oral gavage). Treatment began 16 weeks after tumour initiation by intra-tracheal AdCre delivery. Each bar represents volume change in a single tumour. One-way ANOVA; n.s. P>0.05, * P≤0.05, ** P≤0.01, *** P≤0.001, **** P≥0.0001.
(B) Representative micro-CT scans of mice treated as in (A).

### Supplementary figure 2

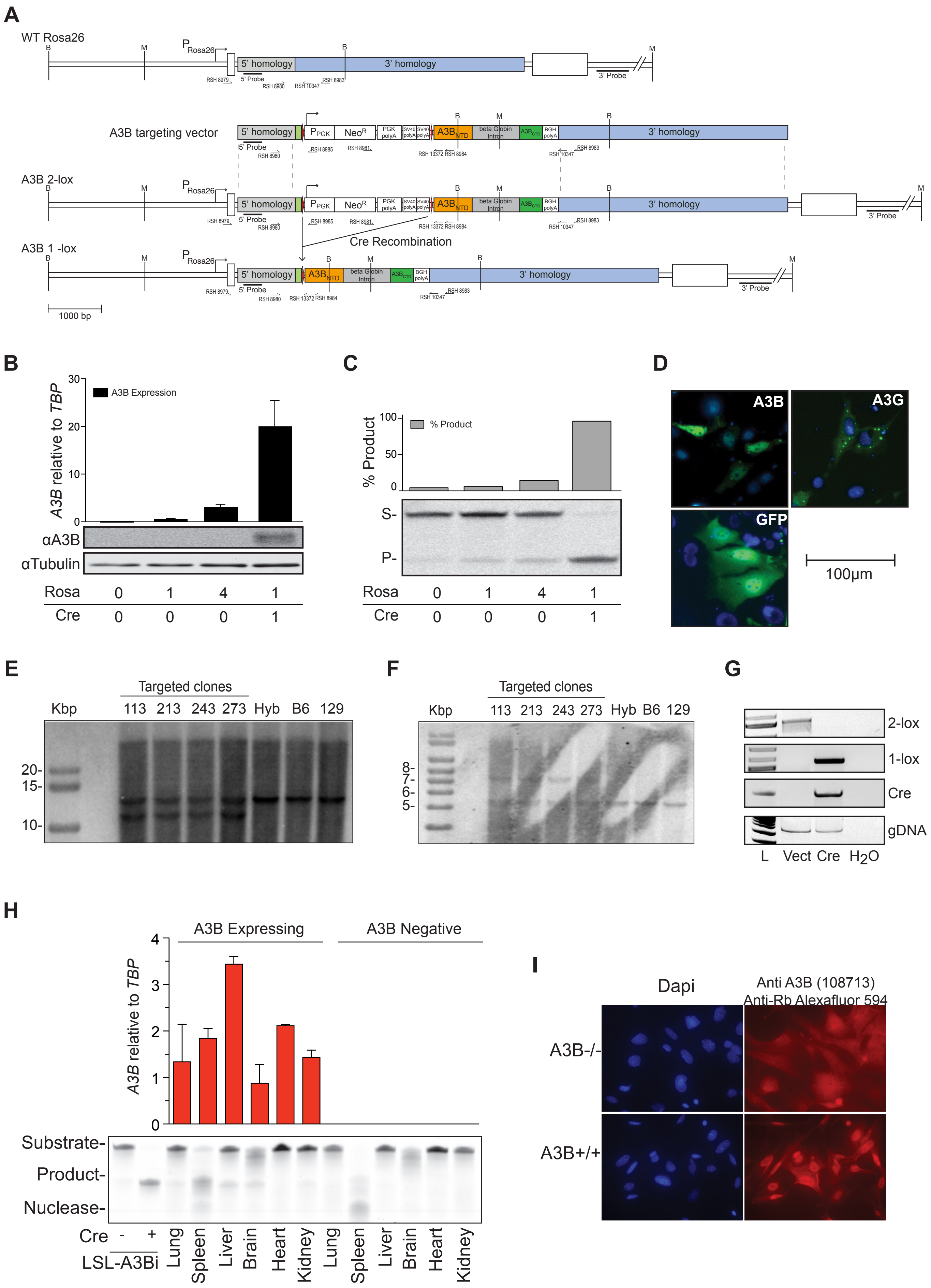

#### Supplementary figure 2 continue

J

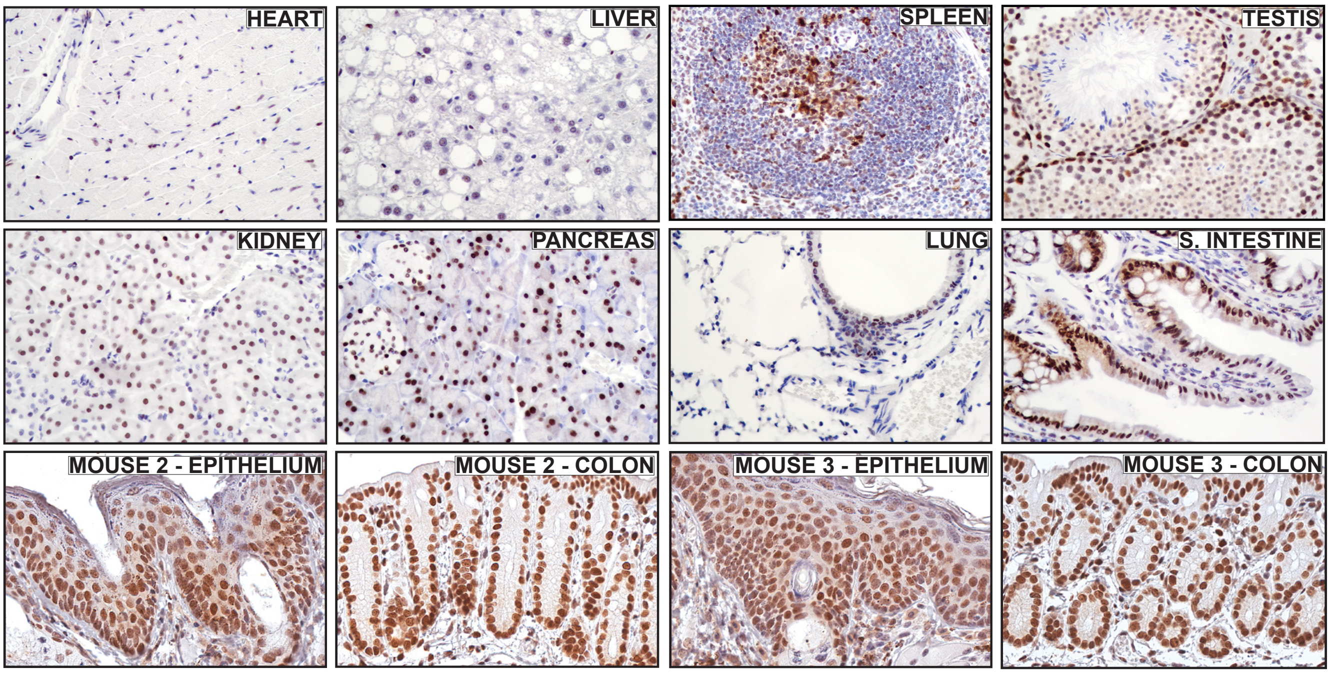


K

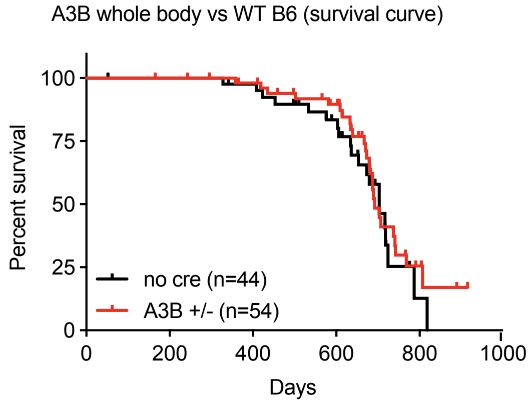


L

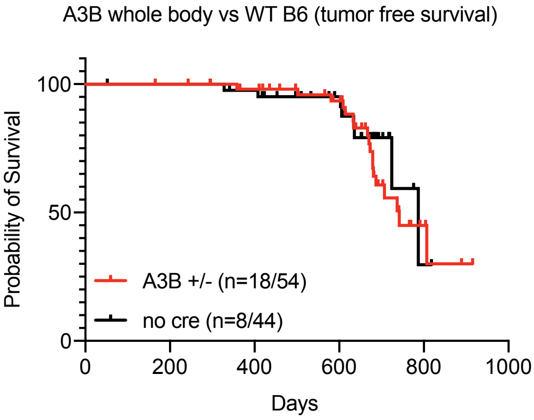


M

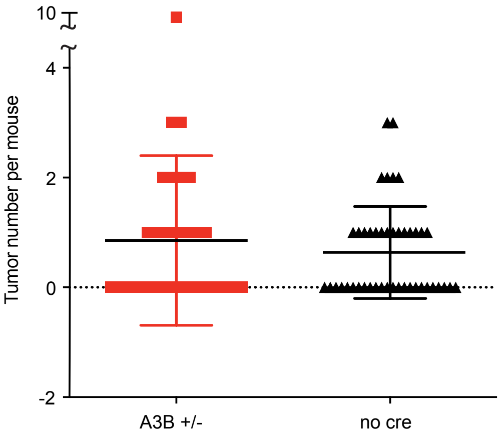

#### Supplementary figure 2 continued

**Supplementary figure 2. Creation and validation of a Cre-inducible human A3B minigene for whole-body or tissue-specific expression in mice (Rosa26::LSL-A3Bi)**

(A) Schematic of the wild-type murine Rosa26 locus, the LSL-A3Bi gene targeting construct with 5' and 3' homology arms, the targeted locus after homologous recombination (Rosa26::LSL-A3Bi), and the targeted locus after Cre-mediated induction of A3Bi minigene expression (Rosa26::L-A3Bi). The chromosomal locus and targeting construct are depicted to-scale with key elements labelled including the loxP-flanked stop cassette (PGK promoter driving a neomycin resistance gene followed by 3 heterologous poly A sites). Relevant restriction sites and primer binding sites are also labelled (M, MfeI; B, BsrG1).

(B) Detection of A3B mRNA (top) and protein (bottom) levels in 293T cells following co-transfection with pcDNA3.1-LSL-A3Bi in the absence or presence of Cre recombinase. The numbers indicate microgram amounts of transfected plasmid.

(C) Quantification (top) of single-stranded DNA C-to-U activity (bottom) in 293T extracts after co-transfection with pcDNA3.1-LSL-A3Bi in the absence or presence of Cre recombinase (S, substrate; P, product). The numbers indicate microgram amounts of transfected plasmid. See panel b for an anti-tubulin control immunoblot.

(D) Fluorescent microscopy images of NIH 3T3 cells transfected with the indicated eGFP-tagged constructs. A3B shows characteristic nuclear localization, A3G is cytoplasmic, and the eGFP control is cell wide. Nuclei are shown in blue (0.01% Hoechst 33342). The scale bar indicates 100 µm.

(E-F) Southern blot analysis of genomic DNA from the indicated PCR-positive ES clones and controls [B6/129 hybrid untargeted ES cells (Hyb), C57BL/6 untargeted ES cells (B6), and SV129 untargeted ES cells (129)]. Genomic DNA was digested with MfeI or BsrGI, fractionated by agarose gel electrophoresis, transferred to membrane, and hybridized respectively with the 3' or the 5' probe depicted in panel a. The MfeI blot yields a 10.96 kbp band for correct targeting and a 12.61 kbp band for the untargeted Rosa26 locus. The BsrGI blot yields a 7.74 kbp band for correct targeting and a 5.61 kbp band for untargeted Rosa26 locus.

(G) Representative results from PCR-based genotyping assays. Non-Cre expressing conditions yield a 500 bp band characteristic of the non-recombined Rosa26::LSL-A3Bi minigene (2x loxP). Cre expression removes the LSL cassette and results in a single loxP site within a characteristic 598 bp PCR product (1x loxP). Specific bands for Cre (100 bp) and murine IL-2 (324 bp) are also shown.

(H) A3B mRNA (top) and DNA C-to-U activity (bottom) levels in the indicated tissues. DNA C-to-U activity in splenic tissue is frequently occluded by nucleolytic degradation of the substrate single-stranded DNA. Activity in heart was not detected due to poor tissue disruption and protein extraction.

(I) Immunofluorescent microscopy of MEFs from control embryos and Rosa26::L-A3Bi expressing embryos. Although the secondary antibody caused a general red fluorescent background, the characteristic nuclear localization A3B is still evident in Rosa26::L-A3Bi expressing embryos.

(J) Immunohistochemical detection of human A3B in the indicated tissues from Rosa26::L-A3Bi mice. The top 8 images are from a representative Rosa26::L-A3Bi male, and the bottom 4 are from two additional animals to demonstrate reproducibility. The characteristic nuclear localization of A3B is clear in all tissues and most cell types. A3B IHC signals are higher in some tissues in comparison to others, likely reflecting natural differences in Rosa26 promoter activity.

(K-L) Kaplan-Meier plots comparing rates of overall survival and fractions of tumour-free animals between a large cohort of Rosa26::L-A3Bi animals (n=54) and a similarly large control cohort with the non-Cre-induced minigene (n=44). Similar numbers of males and females were monitored and no differences were noted between the two sexes.

(M) A dot plot showing the numbers of tumours observed in necropsies of the groups described in panels k-l.

### Supplementary figure 3

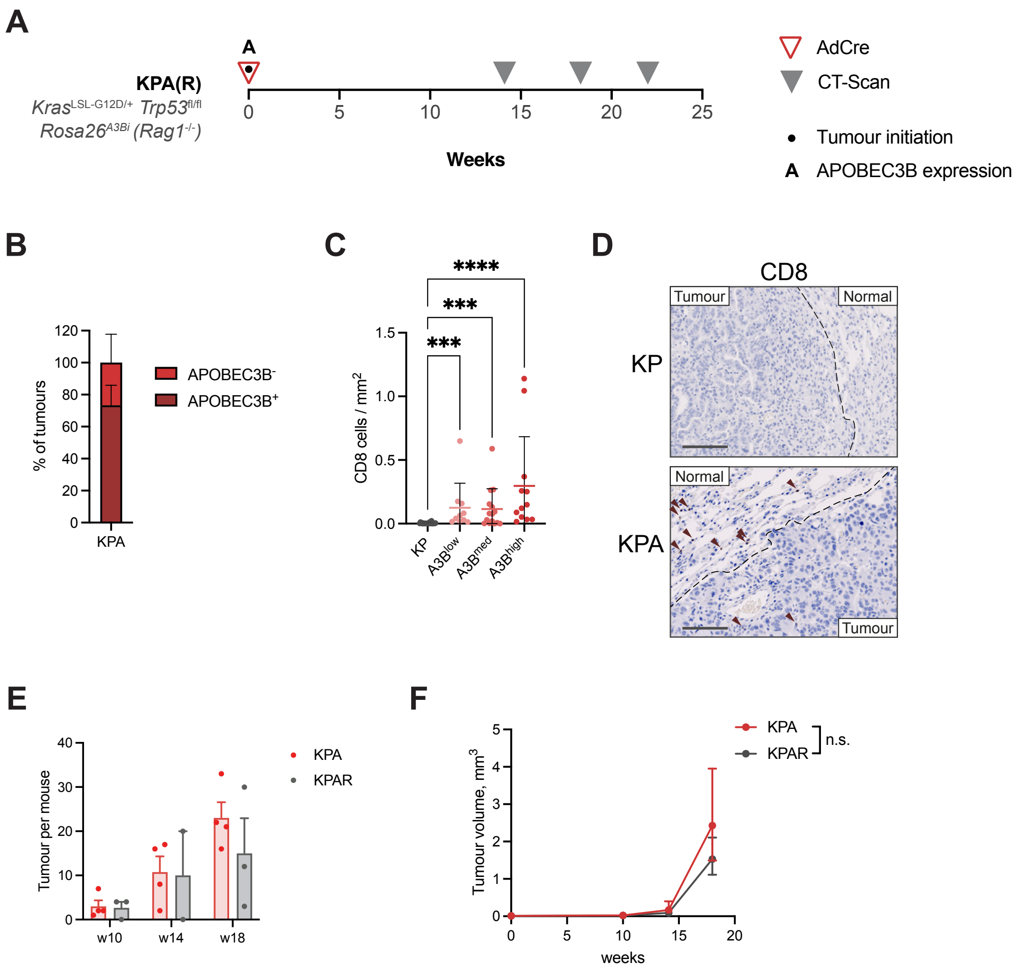


**Supplementary figure 3. A3Bi expression in KP tumours**

(A) Schematic of KPA tumour induction. Tumours were initiated and A3Bi expressed following AdCre (1x106 pfu) intratracheal delivery in KrasG12D/WT; Trp53fl/fl; Rosa26::A3Bi (KPA) or KrasG12D/WT; Trp53fl/fl; Rosa26::A3Bi; Rag1-/- (KPAR) mice. The mice were regularly CT-scanned.

(B) Percentage of APOBEC3B positive and negative tumours in 30 KPA tumour from 5 mice, estimated by immunohistochemistry. Mean, upper and lower limit.

(C) Quantification of immunohistochemistry staining for CD8 in KP and KPA tumours. KPA tumours broken-down according to APOBEC3B expression. Mean per group, ±SD and individual p-value, n=5 mice per group. Kruskal-Wallis test, FDR 0.05; n.s. P>0.05, * P≤0.05, ** P≤0.01, *** P≤0.001, **** P≤0.0001

(D) Immunohistochemistry of CD8 in lung tumours from KP and KPA models. Scale bar represents 100 μm.

(E) Number of tumours per mouse in KPA (n=5 mice) and KPAR (n=3 mice) models estimated from micro-CT scans.

(F) Tumour volume progression in KPA (n=5 mice) and KPAR (n=3 mice) models estimated by micro-CT scans. Mean ±SEM two-way ANOVA, FDR 0.05; n.s. P>0.05.

#### Supplementary figure 4

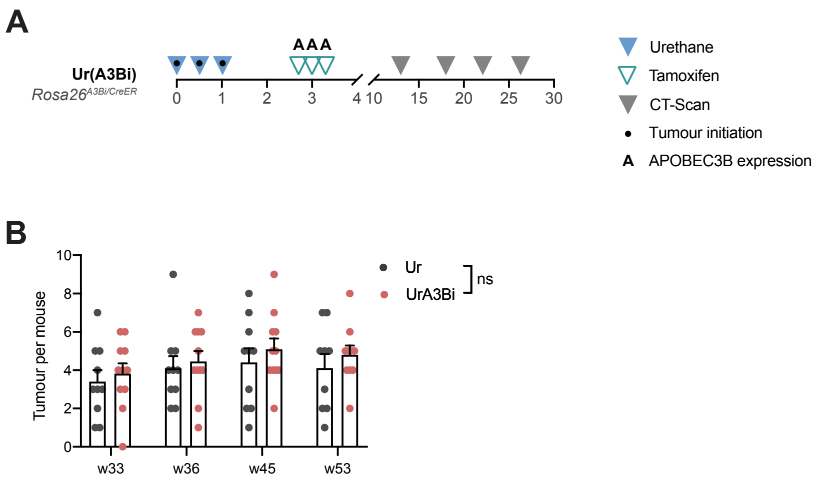


**Supplementary figure 4 | A3Bi expression in urethane-induced tumours**
(A) Schematic of UrA3Bi tumour induction. Tumours were initiated by 3 intra-peritoneal injections of urethane (1 mg/g) and A3Bi expressed following 3 doses of tamoxifen (150 mg/g) (A3Bi-CreER).The mice were regularly CT-scanned.
(B) Mean tumour per mouse in Ur (n=9 mice) and UrA3Bi (CreER) (n=11 mice) models estimated from micro-CT scans. Each coloured dot represents one animal at the indicated time, in weeks after tumour initiation with urethane. Mean expression and SEM. One-way ANOVA, FDR 0.05; n.s. P>0.05

#### Supplementary figure 5

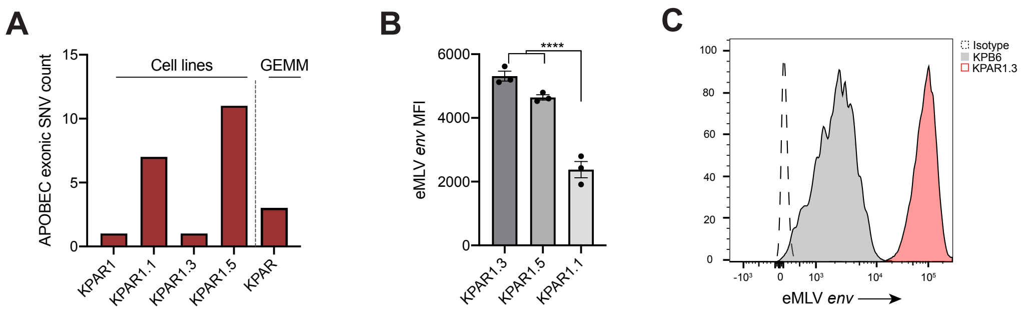


**Supplementary figure 5. Immunogenicity of lung cancer cell lines correlates with eMLV expression**
(A) Frequency of APOBEC-specific (T(C>T/G)) exonic mutations in an autochthonous KPAR tumour, the KPAR parental cell line and the KPAR1.1, KPAR1.3 and KPAR1.5 single-cell clones, estimated by whole-exome sequencing.
(B) Surface expression of eMLV envelope glycoprotein in KPAR1.1, KPAR1.3 and KPAR1.5 cells. Data are mean ± SEM. One-way ANOVA, FDR 0.05; **** P≤0.0001.
(C) Representative histogram plot of eMLV envelope expression on KPAR1.3 and KPB6 cells.

### Supplementary figure 6

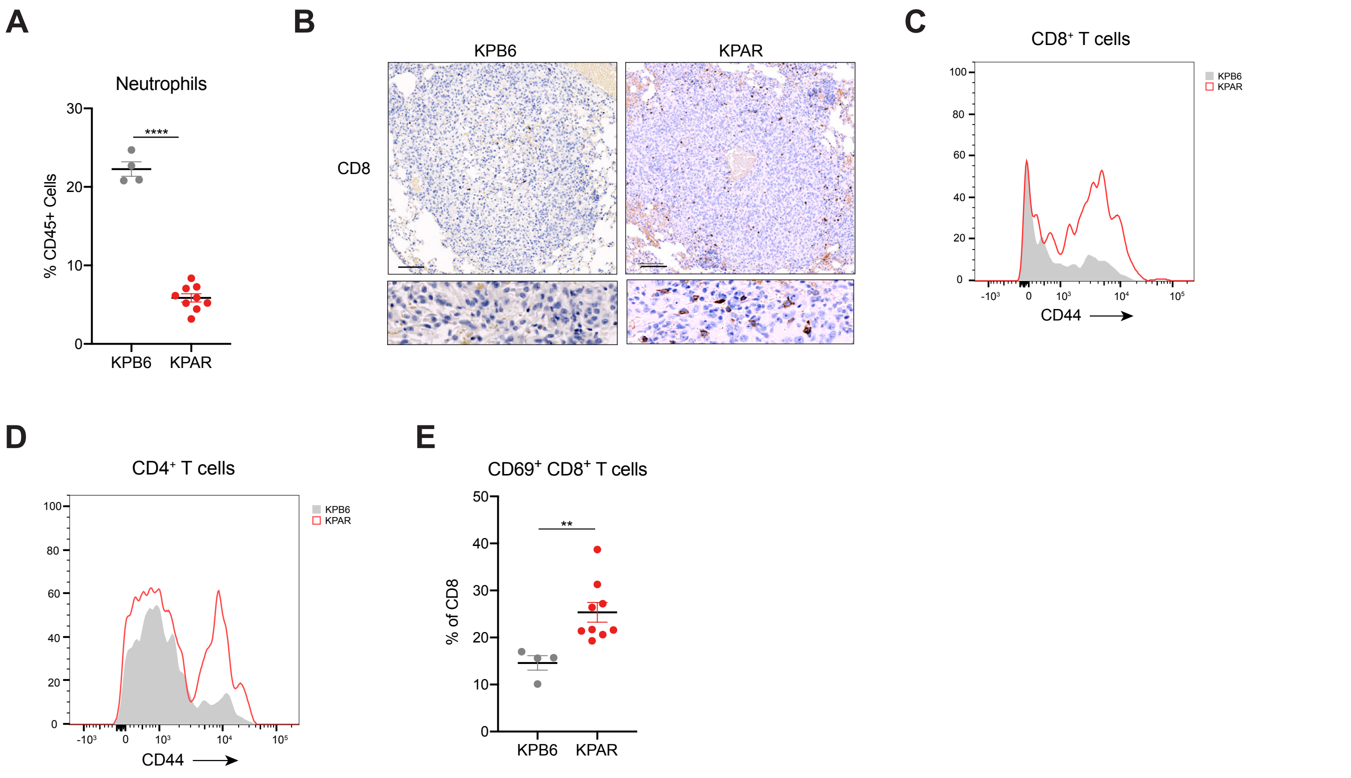


**Supplementary figure 6. Flow cytometry analysis of KPAR and KPB6 orthotopic lung tumours**

(A) Frequency of neutrophils ($CD11b^{+}Ly6G^{+}$) in KPAR and KPB6 orthotopic tumours. Data are mean ± SEM. Unpaired, two-tailed Student's t-test; **** P≤0.0001.

(B) Representative immunohistochemistry staining for CD8 in KPAR and KPB6 orthotopic tumours. Scale bar represents 100 mm.

(C and D) Representative histogram plot of CD44 surface expression on $CD8^{+}$ (C) and $CD4^{+}$ (D) T cells in KPAR and KPB6 orthotopic tumours.

(E) Percentage of $CD69^{+}$ $CD8^{+}$ T cells in KPAR and KPB6 orthotopic tumours. Data are mean ± SEM. Unpaired, two-tailed Student's t-test; ** P≤0.01.

In (A)-(E) tumours were analysed 21 days after transplantation.

### Supplementary figure 7

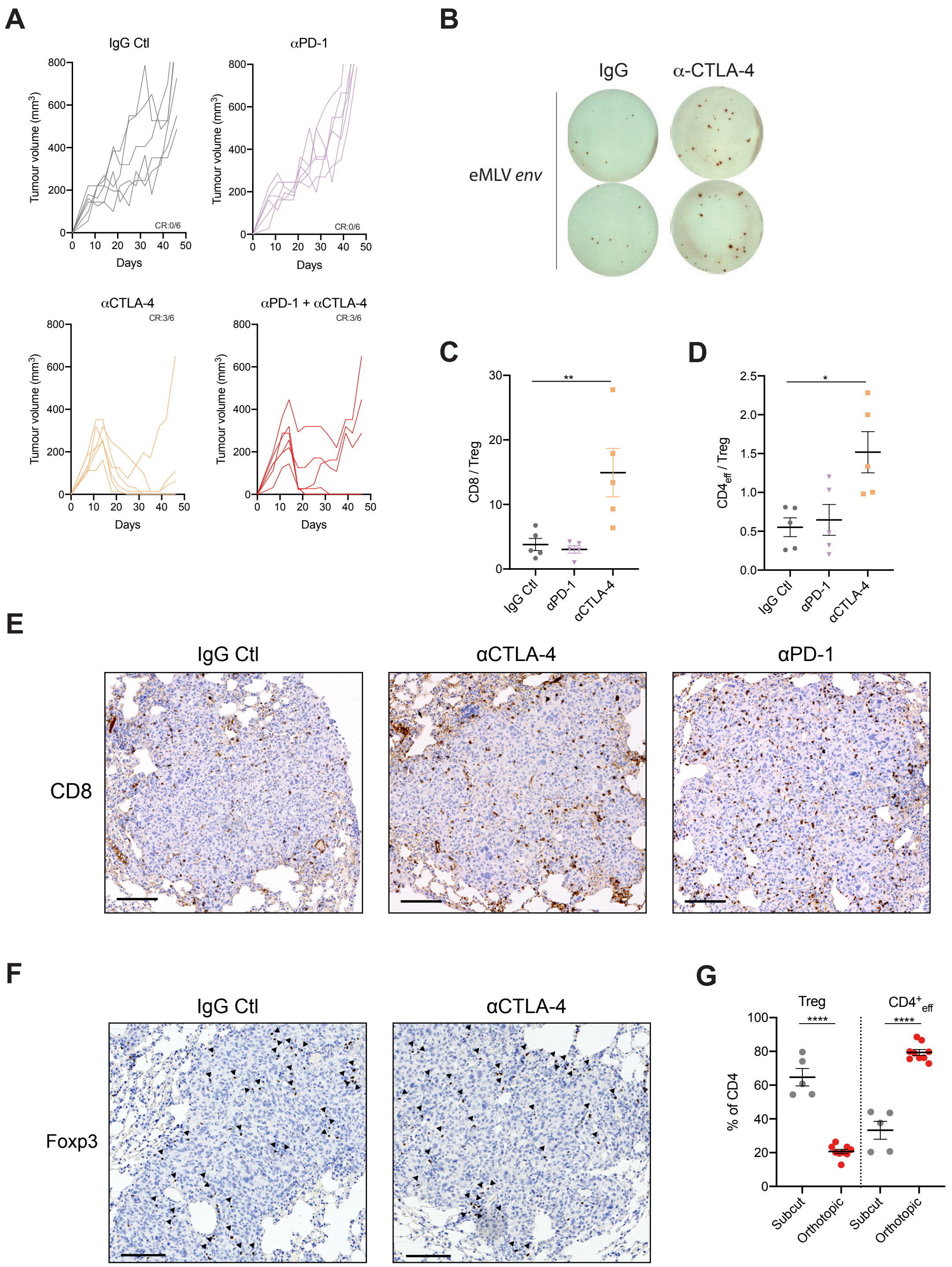


**Supplementary figure 7. KPAR tumours are sensitive to ICB**

(A) Individual subcutaneous KPAR tumour volumes in mice treated intraperitoneally with 200 µg anti-PD-1 and/or 200 µg anti-CTLA-4 or corresponding isotype control on day 10, 14, 17 and 21. CR, complete regression. n=6 mice per group.

(B) IFNγ ELISPOT analysis of TILs isolated from subcutaneous KPAR tumours treated with anti-CTLA-4 or isotype control as in (A). Treatment was on day 10, 14 and 17 and mice were culled on day 18. TILs were pooled from 6 mice per group and pulsed with eMLV *env* peptide.

(C-D) Ratio of $CD8^+$ (C) and $CD4^+$ (D) T cells to $Foxp3^+$ Tregs in subcutaneous tumours treated as in (A). Treatment was on day 10, 14 and 17 and mice were culled on day 18. Data are mean ± SEM, n=5 mice per group. One-way ANOVA, FDR 0.05; * P≤0.05, ** P≤0.01.

(E) Representative immunohistochemistry staining for CD8 in orthotopic KPAR lung tumours treated intraperitoneally with 200 µg anti-PD-1, 200 µg anti-CTLA-4 or corresponding isotype controls twice weekly for two weeks. Treatment was initiated once tumours were detectable by micro-CT. Scale bar represents 100 µm.

(F) Representative immunohistochemistry staining for Foxp3 in orthotopic KPAR lung tumours treated as in (E). Scale bar represents 100 µm.

(G) Frequency of $Foxp3^+$ Tregs and $CD4_{eff}$ ($CD4^+Foxp3^-$) T cells in subcutaneous and orthotopic tumours. Data are mean ± SEM, n=5 mice (subcutaneous) or n=9 mice (orthotopic). Unpaired, two-tailed Student's t-test; **** P≤0.0001.

### Supplementary figure 8

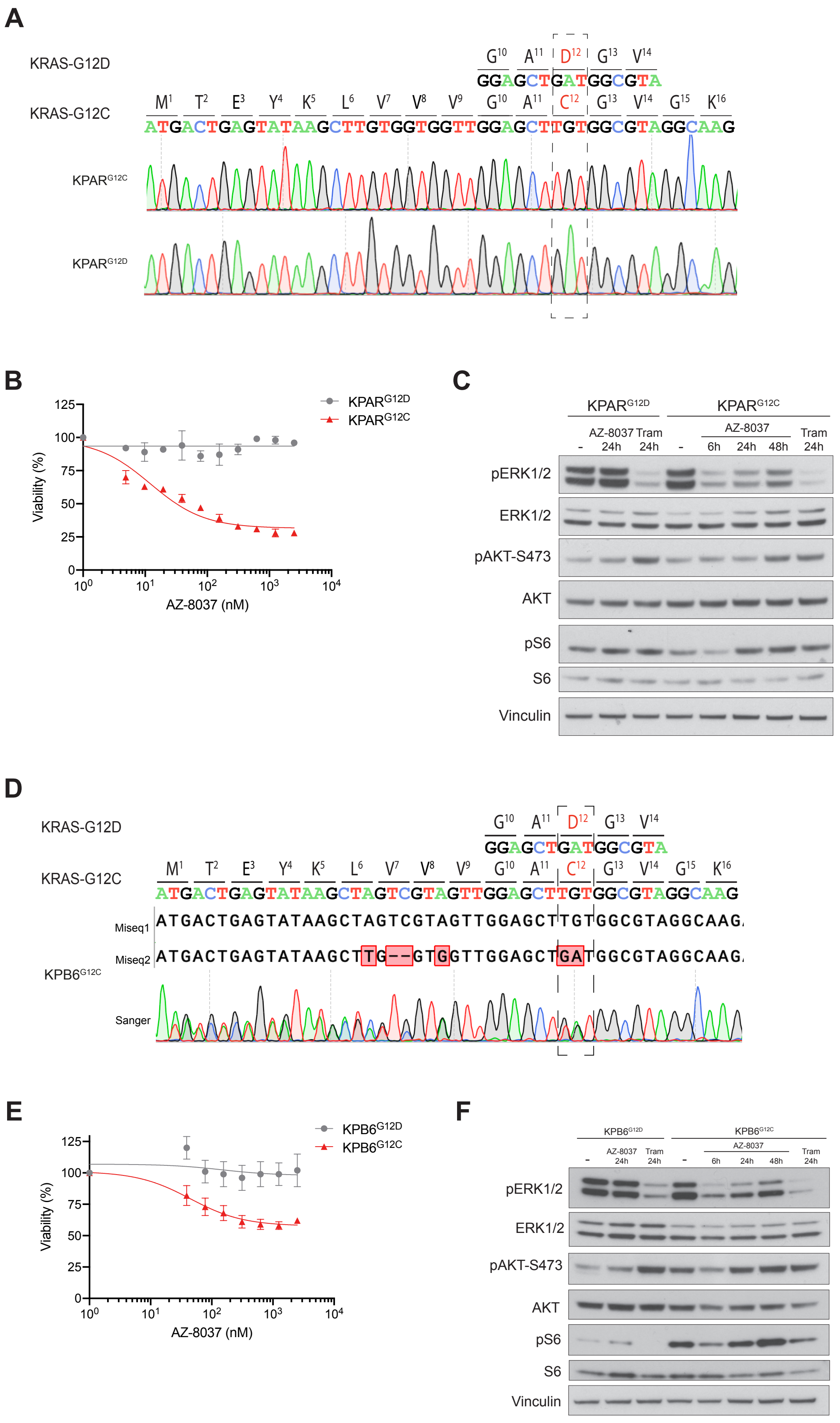

#### Supplementary figure 8 continued

**Supplementary figure 8. Generation of KPAR$^{G12C}$ and KPB6$^{G12C}$ cell lines**

(A) Sanger sequencing chromatograms showing prime-editing of both KRAS$^{G12D}$ alleles of the KPAR cell line into KRAS$^{G12C}$ alleles.

(B) Viability of KPAR parental and KPAR$^{G12C}$ treated with serial dilutions of AZ-8037 for 72h. Data are mean ±SEM of two independent experiments.

(C) Western blot of KPAR parental and KPAR$^{G12C}$ cells treated with 250nM AZ-8037 for 6h, 24h and 48h. Cells were treated with 10nM trametinib (Tram) for 24h as a control.

(D) Sanger sequencing chromatogram and Miseq sequences showing knock-in of the KRAS$^{G12D}$ allele into a KRAS$^{G12C}$ allele (Miseq1) and knock-out of the wildtype allele (Miseq2) in KPB6$^{G12C}$ cells.

(E) Viability of KPB6 parental and KPB6$^{G12C}$ treated with serial dilutions of AZ-8037 for 72h. Data are mean ±SEM of four independent experiments.

(F) Western blot of KPB6 parental and KPB6$^{G12C}$ treated with 250nM AZ-8037 for 6h, 24h and 48h. Cells were treated with 10nM trametinib (Tram) for 24h as a control.
